# Recruitment of TRiC chaperonin in rotavirus viroplasms directly associates with virus replication

**DOI:** 10.1101/2022.12.13.520363

**Authors:** Janine Vetter, Guido Papa, Kurt Tobler, Manuel Kley, Michael Myers, Mahesa Wiesendanger, Elisabeth M. Schraner, Oscar R. Burrone, Cornel Fraefel, Catherine Eichwald

## Abstract

Rotavirus replication takes place in the viroplasms, cytosolic inclusions that allow the synthesis of virus genome segments and their encapsidation in the core shell followed by the addition of the second layer of the virion. The viroplasms are composed of several viral proteins, including NSP5, which is the main building block. Microtubules, lipid droplets, and miRNA-7 are among the host components recruited in viroplasms. To investigate the relationship between rotavirus proteins and host components of the viroplasms, we performed a pull-down assay of lysates from rotavirus-infected cells expressing NSP5-BiolD2. Subsequent tandem mass spectrometry identified all eight subunits of the TRiC complex, a cellular chaperonin responsible for folding at least 10% of the cytosolic proteins. Our validated results show that TRiC is recruited in viroplasms and specifically surrounds newly formed double-layered particles (DLPs). Chemical inhibition of TRiC and silencing of its subunits drastically reduced virus progeny production. Interestingly, TRiC-inhibited RV-infected cells lacked triple-layered particles (TLPs) but harbored empty DLPs. Through sequence-specific direct RNA nanopore sequencing, we show that TRiC is critical for RV replication by controlling dsRNA genome segment synthesis, particularly (-)ssRNA. Moreover, TRiC associates and regulates the folding of VP2, a cofactor allowing dsRNA synthesis. This study provides in-cell culture evidence of the regulatory mechanism by which dsRNA genome segment replication is controlled and coordinated in the rotavirus viroplasms.

**Importance:** The replication of rotavirus takes place in cytosolic inclusions termed viroplasms. In these inclusions, the eleven double-stranded RNA genome segments are synthesized and packaged individually into the newly generated virus particles. In this study, we show for the first time that the TRiC complex, a cellular chaperonin responsible for the folding of at least 10% of the cytosolic proteins, is a component of viroplasms and is required for the synthesis of the viral (-)ssRNA. Specifically, TRiC interacts and assists in folding VP2, the cofactor involved in RNA replication. Our study adds a new component to the current model of rotavirus replication, where TRiC is recruited in viroplasm to assist replication.

## Introduction

Virus factories are compartmentalized inclusions made during the virus life cycle to assemble the replication factors to generate virus progeny. For example, rotavirus (RV), a multisegmented double-stranded RNA (dsRNA) virus and member of the Reoviridae family, replicates in the so-called viroplasms. These structures are composed of several RV proteins and ss- and dsRNAs appearing as cytosolic membrane-less electron-dense inclusions when visualized at the electron microscope (1–3). Several essential processes for the RV life cycle take place in the viroplasms corresponding to the replication and packaging of the eleven dsRNA genome segments (gs) in newly formed VP2 icosahedral core-shells. This process is assisted by replication intermediates composed of the RNA-dependent RNA polymerase (RdRp), VP1, and the helicase and guanylyl-methyltransferase VP3, which are found underneath each of the core-shell five-fold axis (4, 5). VP2 also acts as a cofactor of VP1, at least *in vitro*, for the dsRNA synthesis (6). The nonstructural proteins NSP5 and NSP2 take part in this replication step because they associate with VP1 in a mechanism that is still not completely elucidated (7–9). Subsequently, the filled core shells are coated by a second protein layer of VP6 trimers, forming double-layered particles (DLP), which bud to the adjacent endoplasmic reticulum (ER) to acquire its outer coat. The triple-layered particles (TLP) comprise glycoprotein VP7 and spike protein VP4, which can be found with transient lipid membranes in the ER surrounding the viroplasms (10). Moreover, when detected by fluorescence microscopy, the viroplasms appear as cytosolic globular inclusions that are homogeneously distributed in the cell at early times post-infection (∼4 hpi). The main building block for viroplasm formation is NSP5 (11–13); specifically, its hyperphosphorylated isoform was shown to be essential for forming viroplasms (7, 14–16). Additionally, phosphorylated NSP2 is a requirement for viroplasm formation as well (9, 17). Interestingly, the co-expression of NSP5 with either NSP2 or VP2 is sufficient to build viroplasm-like structures (VLS) (12, 18, 19) which are morphologically identical to viroplasms but lack virus replication components and hence are unable to produce virus progeny. The VLSs are excellent simplified tools for studying the complex viroplasm organization.

It has been demonstrated that viroplasms are highly dynamic, being able to coalesce between them and move to the juxtanuclear region of the cell at increasing times post-infection (20–22). Furthermore, despite not yet being well-defined, several host factors have been identified as necessary for viroplasm formation and maintenance (21, 23–26). On one side, the initiation process for viroplasm formation requires a scaffold of lipid droplets by incorporating perilipin-1 (27, 28). Furthermore, the host cytoskeleton, actin filaments and microtubules, plays a role in the formation, maintenance, and dynamics of the viroplasms (21, 29, 30). In this context, NSP2 octamers directly associate with microtubules to promote viroplasm coalescence (8, 21, 31–33). Moreover, VP2 plays a role in viroplasm dynamics by allowing their perinuclear motion (21). Finally, consistent with the above-described features, the viroplasms have been recently attributed as liquid-liquid phase-separated structures (34).

The RdRp (VP1) shares a unique feature with all described RdRp enzymes of the Reoviridae family (35–38) that consists of four channels that connect the catalytic cavity with the exterior permitting template and NTP entries, and transcript, and template exits (39). This enzyme transcribes from DLPs the mRNA or positive-sense single-stranded RNA ((+)ssRNA) using as a template the negative-sense single stranded RNA ((-)ssRNA) obtained by unwinding the dsRNA genome segments via the C-terminal domains (40, 41). Also, *in vitro* experiments have shown that purified VP1 synthesizes dsRNA using (+)ssRNA as a template by recognizing the 3’ consensus sequence of each genome segment in a process strictly assisted by purified VP2 (6, 42). There is no *in vivo* evidence identifying this mechanism for the encapsidation of the newly generated genome segments.

The eukaryotic group II chaperonin tailless complex polypeptide I ring complex (TRiC) engages in the folding of at least 10% of all cytosolic proteins. It mainly favors the folding of newly translated proteins with complex beta-sheet topologies, such as actin and tubulin, and cell cycle regulators (43). TRiC has a continuously increasing list of client proteins that are involved in diverse cellular processes (44, 45). TRiC is organized as two back-to-back hetero-octameric rings having a barrel-shaped structure enclosing an ATP-dependent folding chamber. Each of these rings comprises eight paralog subunits (CCT1-CCT8) that tweak according to the specificity required for client proteins through differential recognition modes and differentiate rates of ATP binding and hydrolysis between the ring subunits (44, 46). Recently, TRiC activity has also been found to assist the folding of several viral proteins involved in various steps of the virus life cycle, such as entry (47), virus replication (48–54), virion assembly (55, 56) and virus particle release (57).

In this study, we demonstrate that TRiC is essential for RV replication by being recruited to viroplasms, where it assists in dsRNA genome segment synthesis, specifically (-)ssRNA. Furthermore, we reveal that TRiC is associated with NSP5 and VP2 and regulates the folding of VP2, a cofactor for dsRNA synthesis. Finally, we provide *in-cell* culture evidence of the mechanism of RV dsRNA synthesis requires of TRiC-dependent VP2 folding.

## Results

### Association of TRIC components to NSP5-BioID2

To assess the interaction of RV proteins and host components in the viroplasms, we generated a stable MA104 cell line, MA/NSP5-BioID2, expressing NSP5 fused to BioID2, a promiscuous biotin ligase for the detection of protein-protein associations and proximate proteins in living cells (58). As expected (7, 20, 21), NSP5-BioID2 localizes and accumulates upon RV-infection into viroplasms as visualized by the co-localization of streptavidin (StAV)-dylight 488 with VP2, a marker for viroplasms at 6 and 24 hpi **(Fig 1a)**. Next (**Fig 1b**), we analyzed cell extracts of RV-infected MA/NSP5-BioID2 cells at either 5 or 21 hpi by immunoblotting. We found that most biotinylated proteins accumulate in cell extracts prepared at 21 hpi rather than at 5 hpi. Moreover, the bands consistent with the predicted molecular weight of NSP5, NSP2, and VP2 appeared at this specific time upon RV infection. The 21 hpi cell extracts were pulled down with StAv conjugated to magnetic beads and analyzed by label-free tandem mass spectrometry (MS/MS) (**Fig 1c and Table 1**) that enabled the identification of 272 proteins. The distribution of non-infected and RV-infected samples is illustrated in the rank order plot of **Fig 1c**. Consistent with previous publications (14, 20, 29, 59, 60) and validating our experimental procedure, NSP5-BioID2 pulled down RV proteins NSP5, NSP2, VP2, and VP1, which are recognized protein-protein interactors with NSP5. Interestingly, our study also identified the eight subunits (CCT1 to CCT8) of the T-complex protein-1 ring complex (TRiC), a eukaryotic cytosolic ATP-dependent chaperonin that assists the folding of up to 10% of cytosolic proteins (45, 61). To confirm our result **(Fig 1d)**, we pulled down with StAV beads RV-infected MA/NSP5-BioID2 cell extract treated with biotin (added at 1 hpi) and harvested at 6 hpi. As expected, NSP5-BioID2 biotinylated NSP5 and VP2, as denoted by immunoblotting using specific antibodies. We also identified the TRiC subunit CCT1 in both uninfected and RV-infected conditions implicating association with NSP5-BioID2.

**Figure 1.**
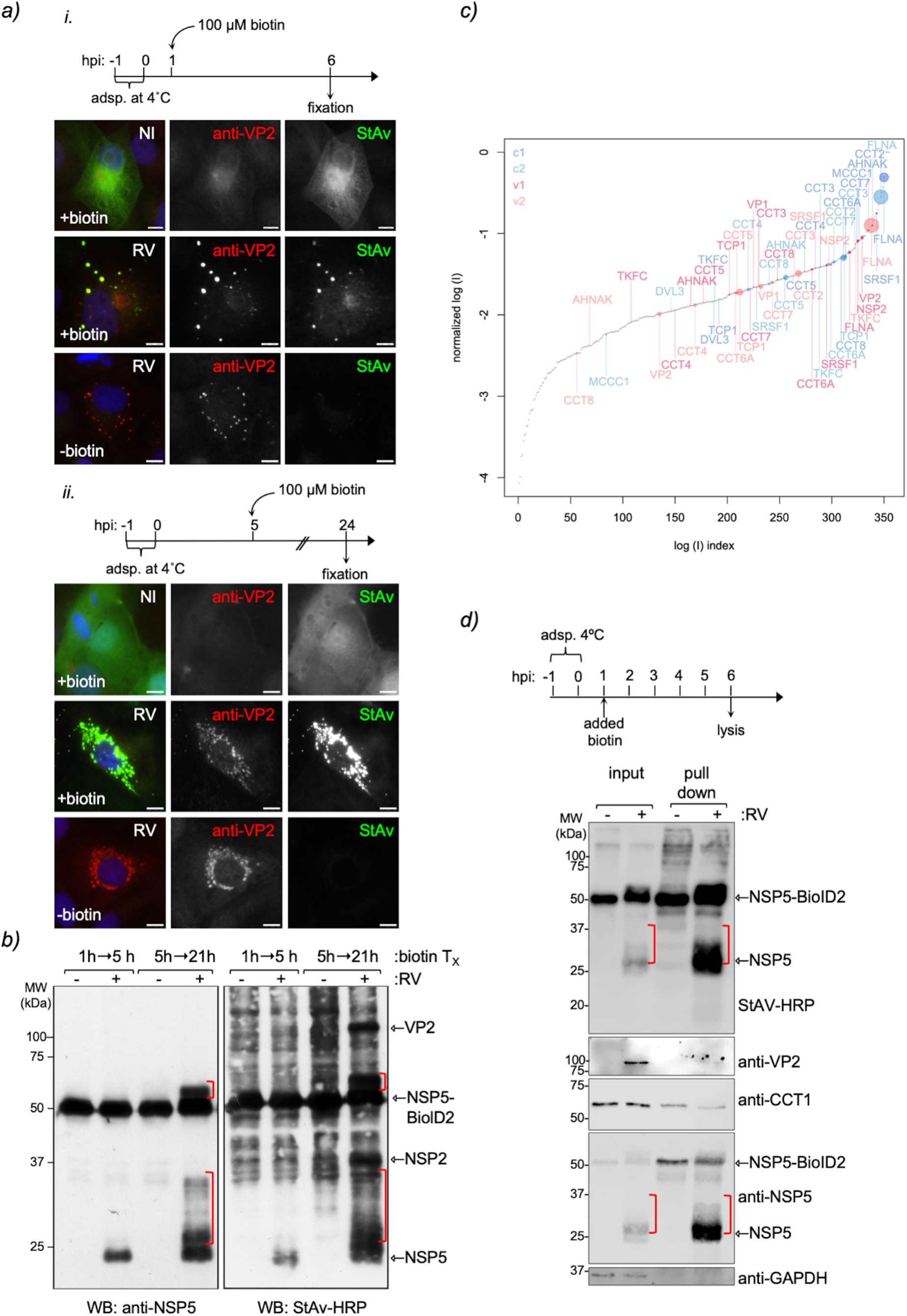
Tandem mass spectrometry analysis of pull-down proteins recruited to NSP5-BioID2 in viroplasms. a) Immunofluorescence images of RV-infected MA/NSP5-BioID2 cells treated with biotin either at 1 hpi (upper panel) or 5 hpi (lower panel) and fixed at 6 or 24 hpi, respectively. Viroplasms were immunostained with anti-VP2 (Alexa 594, red), and NSP5-BioID2 was detected with streptavidin-dylight488 (green). Nuclei were stained with DAPI (blue). Non-infected (NI) control cells are indicated at the top row. The scale bar is 10 µm. **b)** Western blot of non-infected and RV-infected MA/NSP5-BioID2 cell extracts pulled down with streptavidin agarose beads. The cells were treated with 100 µM biotin for the indicated time post-infection (T_x_). The membranes were incubated with anti-NSP5 (left) and streptavidin-HRP (right). Red brackets indicate the NSP5 hyperphosphorylation state. **c)** Mass spectrometry of virus-infected and mock-infected cells. The log(I) values were normalized by subtracting the log(I) value of the NSP5 protein signal. The identified proteins were ranked according to the log of the normalized intensity signal from the mass spectrometry analysis. Proteins identified in the mock-infected samples were colored blue (c1 and c2), and proteins identified in virus-infected samples were colored red (v1 and v2). The sizes of the dots are inversely proportional to the log of the p-values from the mass spectrometry. **d)** Western blot of streptavidin pull-down assay of non-infected and RV-infected NSP5-BioID2 cell lysates. As indicated in the upper scheme, the cells were treated at 1 hpi with biotin and lysed at 6 hpi. The input corresponds to 5% of cell lysates. The membrane was incubated with StAV-HRP and the indicated specific antibodies. Red brackets indicate the NSP5 hyperphosphorylation state.

**Table 1.**
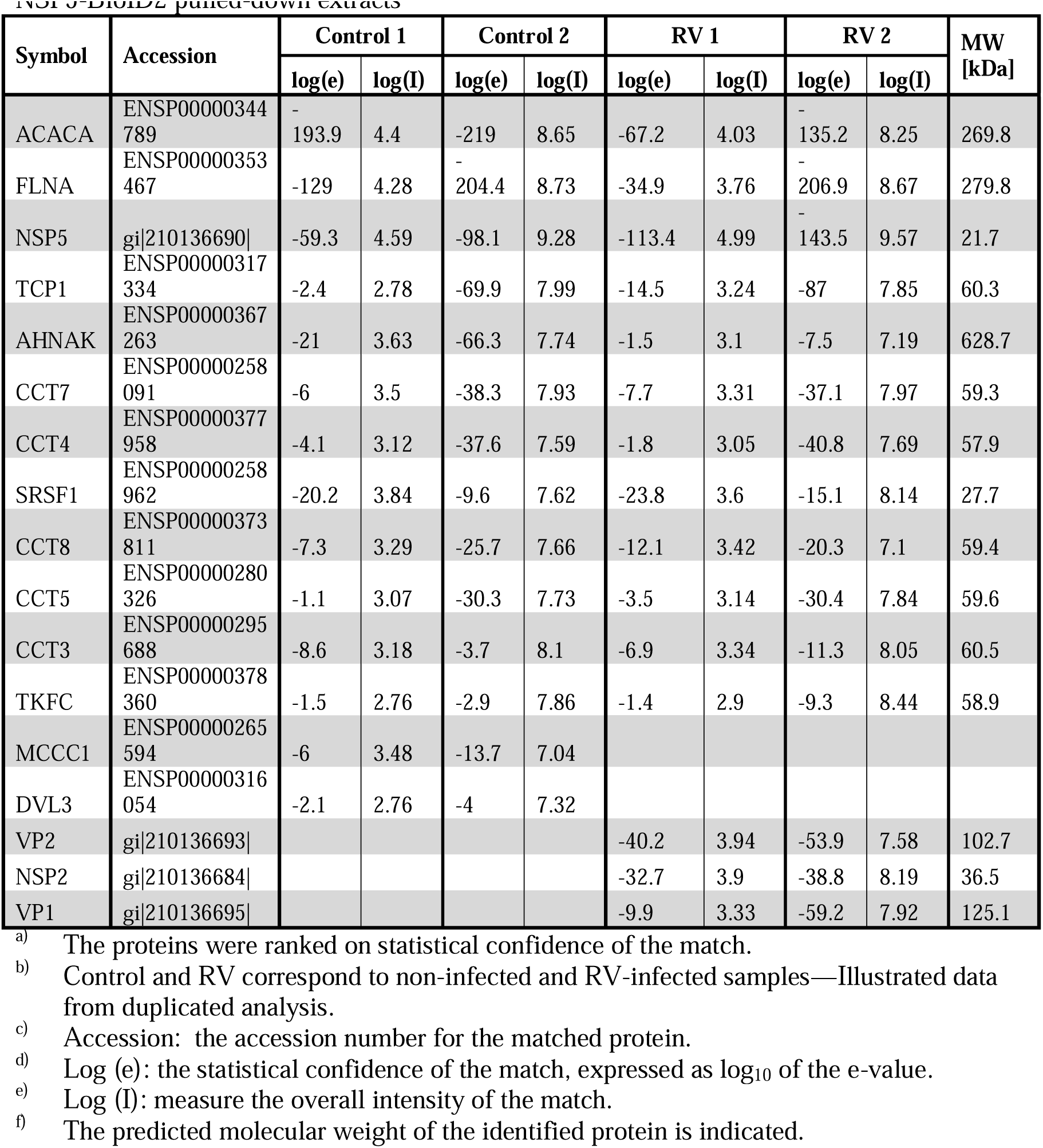
List of ranked data obtained from mass spectrometry analysis of RV-infected MA-NSP5-BioID2 pulled-down extracts.

### TRiC localizes in viroplasms and surrounding DLPs

We next investigated whether TRiC localizes into viroplasms at 6 hpi to validate the mass spectrometry analysis. As visualized with immunofluorescence confocal microscopy (**Fig 2a)**, TRiC subunits CCT1, CCT2, and CCT3 colocalize with NSP5, a viroplasm marker, suggesting that TRiC is recruited to viroplasms. No crossreactivity was observed between TRiC-specific antibodies and RV antigens (**Fig S1**). Next, we also analyzed viroplasms at high resolution using immune electron microscopy by co-immunostaining of anti-CCT3 (12 nm gold particles) with either anti-NSP5 (6 nm gold particles, **Fig 2b**) or anti-VP6 (6 nm gold particles, **Fig 2c**). Interestingly, CCT3, in addition to localizing in viroplasms, was found in some cases surrounding structures resembling DLPs by their morphology and size and because VP6 surrounds them. Interestingly, NSP5 and CCT3 are also found to encompass these globular structures. Like CCT3, also CCT2 colocalizes in viroplasms circumscribing DLPs (**Fig S2**).

**Figure 2.**
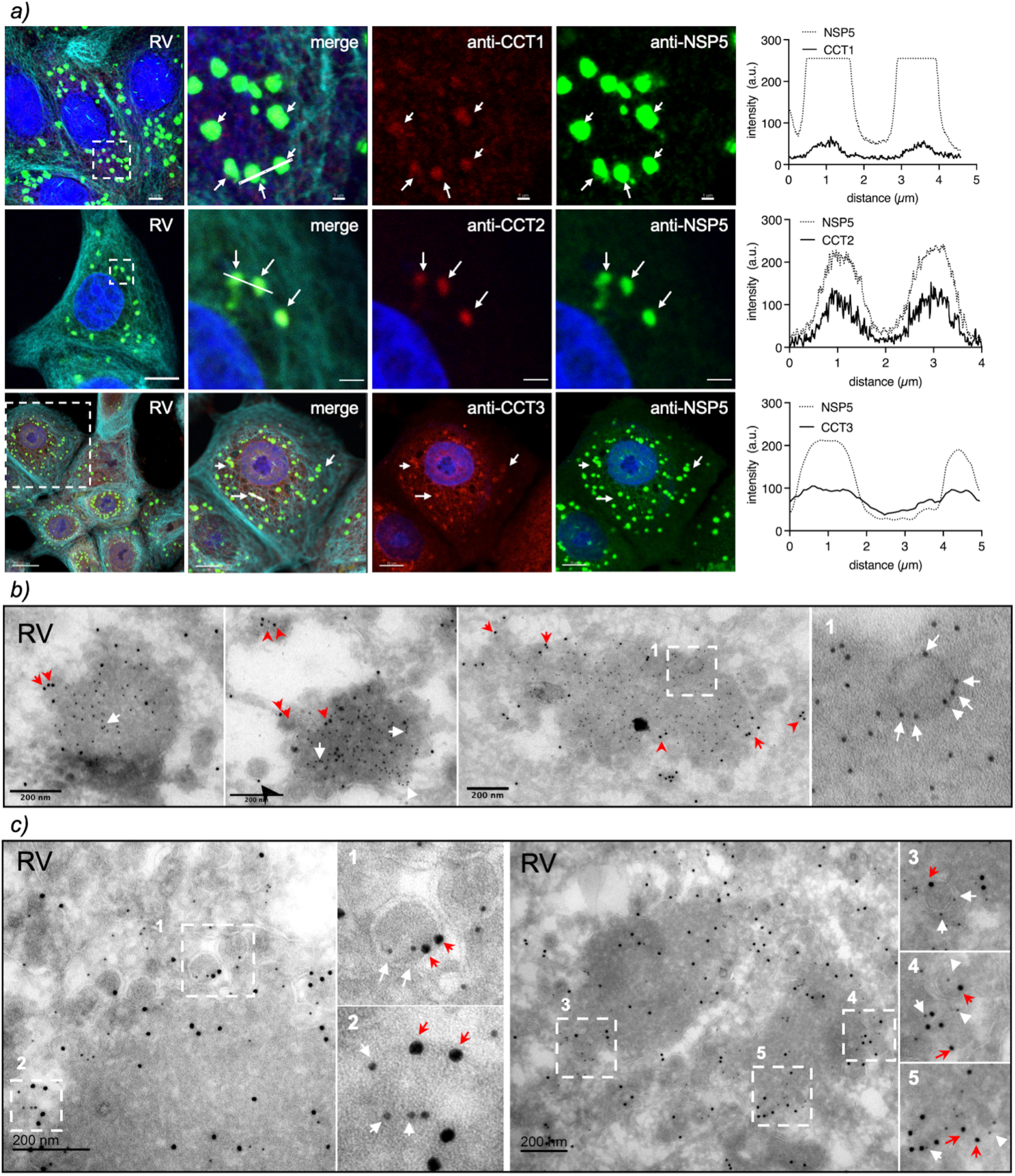
TRiC subunits localize in viroplasms surrounding virus particles. a) Immunofluorescence of RV-infected cells immunostained at 6 hpi for the detection of viroplasms (anti-NSP5, Alexa 488, green), microtubules (anti-alpha tubulin, Alexa 647, cyan), and TRiC subunits CCT1, CCT2, and CCT3 (Alexa 594, red). Nuclei were stained with DAPI (blue). The white dashed box represents the enlarged image at the right. White arrows point to the co-localization of viroplasms with TRiC subunits. The scale bar is 10 µm. The plots in the right column correspond to the co-localization profile of the linear region of interest of NSP5 with the TRiC subunit. Immune electron microscopy of viroplasm fixed at 6 hpi. The thin sections were co-immunostained with either anti-NSP5 conjugated to 6 nm gold **(b)** or anti-VP6 conjugated to 6 nm gold **(c)** followed by anti-CCT3 conjugated to 12 nm gold. The white dashed open boxes correspond to enlarged indicated images. Red arrowheads and white arrows point to the localization of CCT3 and NSP5 or VP6 surrounding DLPs. The scale bar is 200 nm.

### Inhibition of TRiC hampers viroplasm formation and RV replication

We used a recently described chemical TRiC inhibitor (62) (PubChem CID: 658022) corresponding to 2-[(4-chloro-2λ4,1,3-benzothiadiazol-5-yl)oxy]acetic acid, shortly named TRICi, to investigate the role of TRiC chaperonin in the RV life cycle. TRICi was validated in MA104 cells for its ability to halt the onset of mitosis by impairing Cdc20 expression **(Fig S3a-e),** a well-described protein dependent on TRiC for folding (63). When inspecting viroplasms at 6 hpi, a time-frame for visualization of well-formed viroplasms (21), RV-infected cells treated at early time post-infection (1 hpi) with TRICi showed significantly reduced numbers and size of viroplasms per cell (**Fig 3a, b, and c**). In contrast, the addition of TRICi at 5 hpi showed no changes in the morphology and number of viroplasms compared to untreated samples (**Fig 3a**). The TRICi treatment was not cytotoxic at the concentrations used in our experiments **(Fig S3f)**. Notably, the impairment in viroplasm formation after TRiC inhibition was observed for at least three RV strains, including porcine strain OSU and simian strains SA11 and RRV (**Fig S4**). The virus progeny was largely impaired (> 5 log) when TRICi was added at 1 hpi at both tested concentrations but not when added at 5 hpi **(Fig 3d)**. Similarly, silencing either CCT3 or CCT2 subunits significantly reduced virus progeny **(Fig S5a-d)**. Consistent with the lack of virus progeny, we observed a complete depletion of the RV dsRNA genome segments (**Fig 3e**) after TRICi treatment (lanes 2 and 3) when compared to untreated cells (lane 1).

**Figure 3.**
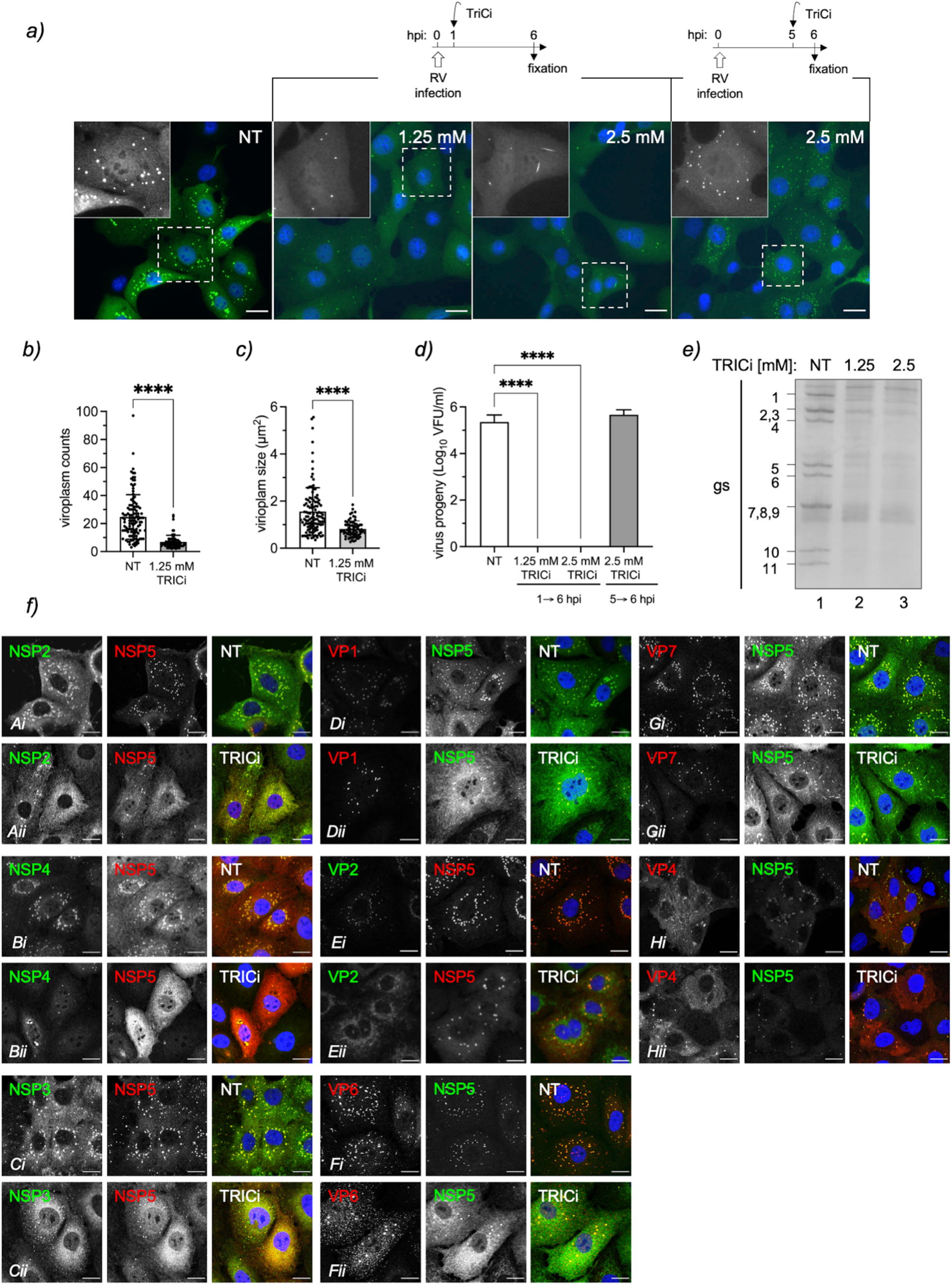
TRiC inhibition impairs viroplasm morphology and virus progeny. a) Immunofluorescence micrograph of OSU-infected MA104 cells untreated or treated with 1.25 mM or 2.5 mM TRICi and fixed at 6 hpi. The drug inhibitor was added at 1 or 5 hpi as indicated. Cells were immunostained to detect viroplasms (anti-NSP5, green). Nuclei were stained with DAPI (blue). An enlarged image of a single cell is provided in black and white. The scale bar is 10 µm. Plots for quantification of numbers **(b)** and size **(c)** of viroplasms per cell after TRICi treatment from 1 to 6 hpi. Data represent the mean± SD. n> 50 cells; (****), p<0.0001. **d)** Plot for virus progeny of RV-infected cells treated with TRICi during the indicated time post-infection. Data represent the mean± SD of three independent experiments; (****), p<0.0001. **e)** Electropherotype of RV genome segments (gs) extracted at 6 hpi from RV-infected cells non-treated (NT) and treated with 1.25 mM or 2.5 mM TRICi for 5 h before cell lysis. **f)** Immunofluorescence micrograph of OSU-infected cells showing the distribution of RV proteins after treatment with 1.25 mM TRICi since 1 hpi. At 6 hpi, cells were fixed and immunostained to detect viroplasms (anti-NSP5; guinea pig polyclonal (green) or mouse monoclonal (red) antibodies) and the indicated RV protein (using specific antibodies for each of them). Nuclei were stained with DAPI (blue). The capital letters in the lower left corner correlate with the analyzed protein: A, NSP2; B, NSP4; C, NSP3; D, VP1; E, VP2; F, VP6; G, VP7; H, VP4. Each panel shows untreated (*i*, NT) and 1.25 mM TRiC treated (*ii*, TRICi) samples. The scale bar is 10 µm.

Additionally, the effect of TRICi on viroplasms was partially reversible when added at 1 hpi and washed out at 2, 3, 4, or 5 hpi (**Fig S6a-e**), followed by analysis at 6 hpi. Large viroplasms recovered even when removing the drug at 4 hpi, reaching the same levels as untreated samples. Meanwhile, the number of small viroplasms did not recover even after only 1 h of treatment with TRICi, remaining in large numbers distributed in the cytosol of the infected cells. Besides, the expression of diverse RV proteins was recovered after removing TRICi **(Fig S6f)**, which collectively suggests a delay in the coalescence of viroplasms because of a lack of RV protein supplies required for building the inclusions. Additionally, the distribution of diverse RV proteins was compared in untreated and TRICi-treated conditions by immunofluorescence at 6 hpi. As observed in **Fig 3f**, RV viroplasm proteins NSP5, NSP2, VP2, and VP6 delocalized from the viral factories and dispersed throughout the cytosol, while VP1, NSP4, VP7, and VP4 did not show perceptible changes from their characteristic distribution. Also, NSP3 showed dramatic changes with a dispersed distribution.

### TRiC inhibition hampered RV packaging and TLP formation

To investigate in detail the decrease in both dsRNA genome segments and virus progeny associated with TRiC inhibition, we examined the viroplasms by high-resolution electron microscopy at 6 hpi after treatment with TRICi from 1 hpi. Surprisingly **(Fig 4a)**, we observed that both untreated and TRICi-treated viroplasms were electron-dense. As expected, the endoplasmic reticulum (ER) surrounding the viroplasms in the untreated sample was filled with TLPs at diverse stages of maturation. Meanwhile, TRICi-treated samples showed almost empty ER with a few immature virus particles. We then analyzed the virus particles isolated at 24 hpi from cells treated with TRICi at 1 hpi or untreated to gain insights into virion structure and composition. We found that the negatively stained virus particles from TRICi-treated cells had an average size of 55 nm, consistent with the size and shape of DLPs. In contrast, the virus particles from untreated cells had an average size of 76 nm, consistent with the size of TLPs **(Fig 4b)**. Moreover, we compared untreated and TRiC-treated RV-infected cell extracts harvested at 8 hpi and found that VP7 was significantly reduced (**Fig 4c and d**). Interestingly, an equivalent amount of purified virus particles isolated from TRICi-treated samples lacked the outer layer proteins VP7 and VP4 but also had reduced amounts of VP1 and low association with NSP2 (64) **(Fig 4c, e, and f**). This outcome contrasts with the virus particles from untreated cells containing all TLP components. Upon these conditions, the particles from TRICi-treated cells also had significantly reduced dsRNA genome segments with an average of approximately 60% less virus genome per DLPs than untreated samples **(Fig 4g, h, and i)**. This outcome suggests a preponderance of DLPs over TLPs, with accumulation of empty particles lacking the virus genome upon TRiC inhibition.

**Figure 4.**
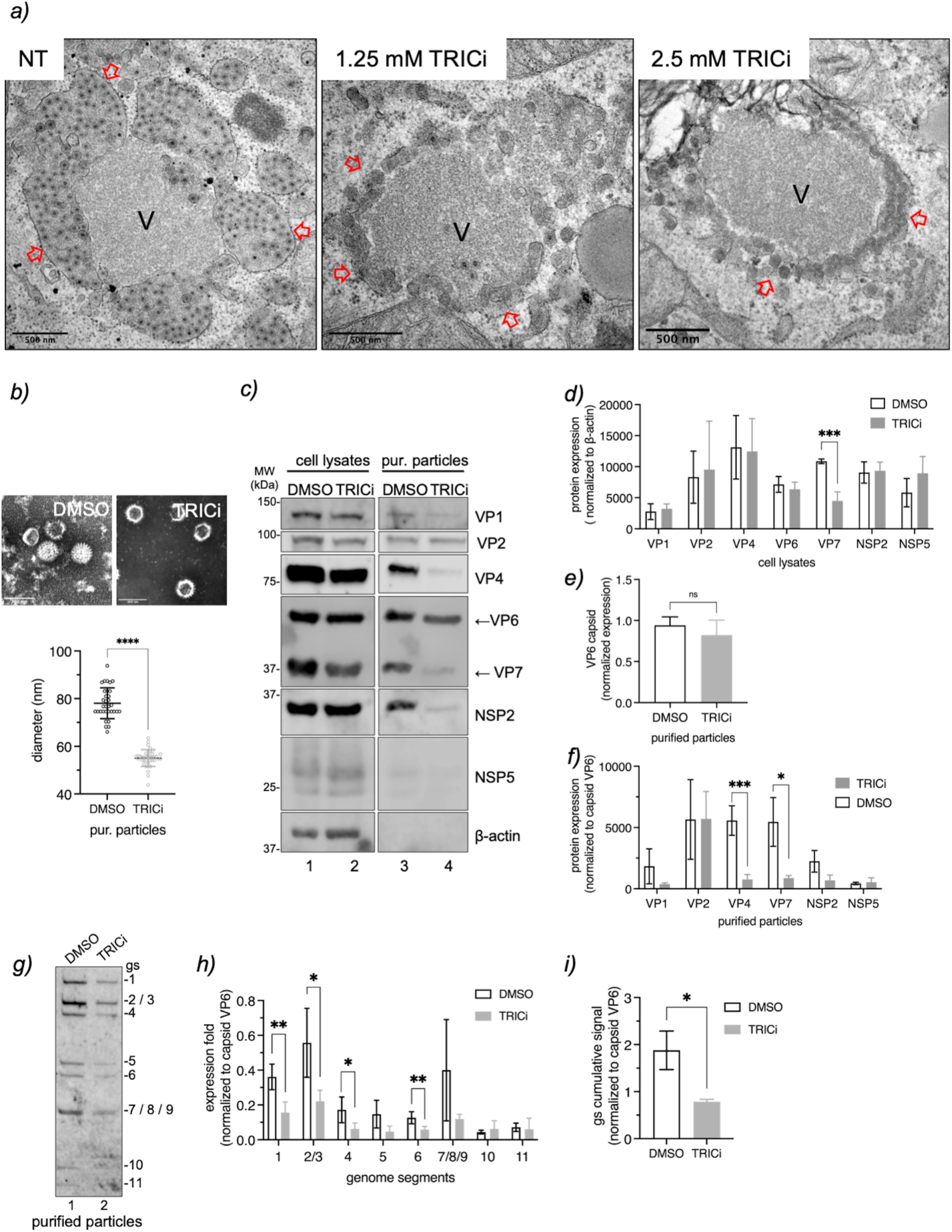
Inhibition of TRiC leads to empty DLPs. a) High-definition electron microscopy of OSU-infected MA104 cells untreated and treated with TRICi at the indicated concentrations. The inhibitor was added at 1 hpi, and the samples were fixed at 6 hpi. The red, open arrowheads point to the endoplasmic reticulum surrounding viroplasms. The scale bar is 500 nm. **b)** Negative staining of purified virus particles isolated from OSU-infected cells untreated or treated with 1.25 mM TRICi from 1 hpi. The scale bar is 100 nm. The plot corresponds to the size mean ± SEM of purified particles after indicated treatments. (***), p-value < 0.001. **c)** Immunoblot of OSU infected MA104 cell lysates (left panel) and the purified virus particles (right panel) at 8 hpi after treatment without (lanes 1 and 3) or with 1.25 mM TRICi (lanes 2 and 4) added at 1 hpi. The membranes were incubated with the indicated specific antibodies. d) Plot comparing the expression of RV proteins in cell lysates untreated or treated with TRICi. The samples were normalized to housekeeping protein beta-actin. **e)** Plot of comparing the expression of middle layer capsid proteinVP6 of purified virus particles. **f)** Plot comparing the expression of RV proteins present in purified virus particles normalized to VP6 capsid expression. The plots represent the mean ± SD of at least three independent experiments. **g)** Electropherotype of OSU genome segments extracted from purified virus particles of cells treated with 1.25 mM TRICi from 1 hpi to 8 hpi. The corresponding dsRNA genome segments are indicated. Plots comparing normalized expression fold of each OSU genome segment **(h)** or their cumulative signal **(i)** when extracted from purified particles untreated or treated with 1.25 mM TRICi. The bands corresponding to genome segments 2/3 and 7/8/9 were assessed as unique values, respectively. The data correspond to the mean ± SD of three independent experiments, two-way ANOVA where (*), p<0.05 and (**), p<0.01.

### Synthesis of (-)ssRNA is reliant on TRiC

We next hypothesized that TRiC inhibition blocks the synthesis of (-)ssRNA but not (+)ssRNA. To assess this possibility, we established a method for direct RNA sequencing for RV dsRNA to read both positive- and negative-sense RNA strands using MinION Oxford nanopore technology (ONT) (65). For this purpose, we designed specific reverse transcriptase adapters (RTA) that anneal to the 3’ ends of both positive- and negative-sense RNA strands of the eleven genome segments of porcine RV strain OSU **(Table S2)**. Total RNA from OSU-infected cells at 6 hpi was harvested and sequenced via MinION to determine the effectiveness of the modified adapter. The sequence runs covered 100% of the eleven genome segments of both positive- and negative-sense RNA strands **(Fig 5a)**. The average coverage depth for the positive-sense RNA strands ranked from 7987.09 for gs 8 (NSP2) to 528.93 for gs5 (NSP1), while the negative-sense RNA strands ranked from 1632.10 for gs 11 (NSP5) to 132.48 for gs 4 (VP4), respectively **(Table 2)**. The average read level of accuracy of the eleven genome segments was 91.28 ± 0.46 % and 90.05± 0.58% for the positive- and negative-sense RNA strands, respectively. Meanwhile, the sequence coverage corresponded to 99.47±0.32% for the positive-sense RNA strands and 99.28±0.41% for negative-sense RNA strands, in concordance with the consensus sequence of the eleven genome segments provided for the RV strain OSU. The distribution of positive- and negative-sense RNA strands read lengths **(Fig 5b)** correspond well to the expected length of each respective segment. An odd cumulative ratio **(Fig 5c)** was determined between the distribution of positive and negative sense RNA strand reads for each genome segment with an accumulative value of 453. We then harvested RNA at 6 hpi from RV-infected cells either untreated (DMSO) or treated with TRICi and followed direct RNA sequencing of both positive- and negative-sense RNA strands. The sample from DMSO-treated cells showed a complete coverage for both positive- and negative-sense RNA strands (**Fig 5d, upper panel; Table 3**), with a similar distribution pattern observed for the above mock sample (**Fig 5b, e, and f)**. However, the samples from TRICi-treated cells, although showing similar depth coverage for all the eleven positive-sense strands as for DMSO samples, denoted a highly reduced coverage for the eleven negative-sense strand RNAs **(Fig 5, lower panel)**. Even if the distribution of the eleven genome segments for positive-sense RNA strand reads **(Fig 5e)** was comparable between samples from untreated and TRICi-treated cells, it was highly diverse for the negative-sense reads **(Fig 5f)**. More precisely, the numbers of negative-sense RNA strand reads were at least three times lower in the TRICi-treated samples compared to the untreated samples. The ratio of positive-/negative-sense RNA strand reads **(Fig 5g)** for DMSO-treated cells had a value of 509, similar to the above-described value, while that of the TRICi-treated cells was much higher with a value of 1897, suggesting a preponderance of (+)ssRNA reads among the TRICi sample.

**Figure 5.**
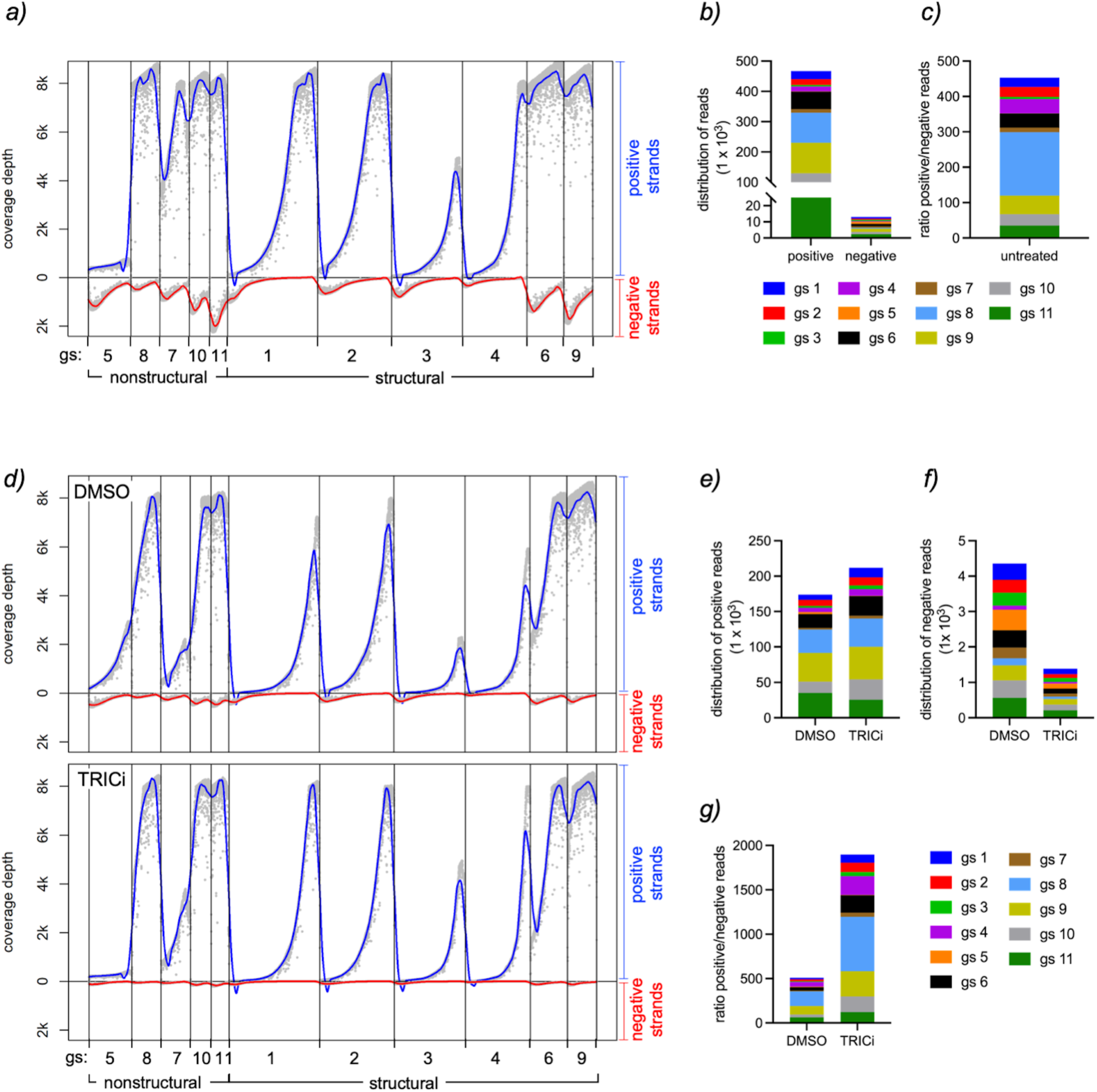
Rotavirus negative-strand synthesis is hampered upon inhibition of TRiC. a) Oxford nanopore technology (ONT) for direct sequencing of rotavirus positive and negative sense strand RNA of the eleven genome segments at 6 hpi. The plot indicates the depth of the sequence coverage of the positive (blue line) and negative (red line) sense strand RNA for the eleven RV genome segments. The genome segments encoding for nonstructural and structural proteins are indicated. **b)** Plot indicating the distribution of positive and negative sequence reads for each of the eleven genome segments. **c)** Plot indicating the cumulative ratio between positive and negative RNA sequence reads for the eleven genome segments. **d)** Direct RNA sequencing using ONT of RV-infected cell extracts untreated (DMSO, upper panel) or treated with 2.5 mM TRICi (TRICi, lower panel) at 6 hpi. The chemical compound was added at 1 hpi until cell lysis. The plot indicates the depth of the sequence coverage of the positive (blue line) and negative (red line) for the eleven RV genome segments. The genome segments encoding for nonstructural and structural proteins are indicated. Plot for the distribution of the positive **(e)** and negative **(f)** sequence reads of RNA isolated from DMSO and TRICi-treated RV-infected cells. **(g)** Plot comparing the ratio between positive and negative RNA sequence reads of RV-infected cell extracts treated with DMSO or TRICi.

**Table 2:** ONT average coverage depth for (+) and (-) ssRNAs isolated from cells infected with porcine OSU strain.

| RNA sequence<sup>a)</sup> | Average coverage depth (reads/nucleotide) |  |
| --- | --- | --- |
|  | positive strand | negative strand |
| gs 1 (VP1) | 3282.45 | 196.16 |
| gs 2 (VP2) | 3901.18 | 261.65 |
| gs 3 (VP3) | 1223.30 | 271.19 |
| gs 4 (VP4) | 2109.55 | 132.48 |
| gs 5 (NSP1) | 528.93 | 671.66 |
| gs 6 (VP6) | 7705.13 | 864.61 |
| gs 7 (NSP3) | 7465.08 | 607.89 |
| gs 8 (NSP2) | 7987.08 | 343.86 |
| gs 9 (VP7) | 5845.46 | 1154.66 |
| gs 10 (NSP4) | 7906.21 | 1060.12 |
| gs 11 (NSP5) | 7858.12 | 1632.10 |
| <sup>b)</sup> GAPDH | 4.12 | 0.00 |

<sup>a)</sup> Rotavirus porcine strain OSU.
<sup>b)</sup> Sequence covering GAPDH of *Macaca fascicularis*

**Table 3:** ONT average coverage depth for (+) and (-)ssRNAs isolated of OSU infected cells untreated or treated with TRICi.

| RNA sequence<sup>a)</sup> | Average coverage depth (reads/nucleotide) |  |  |  |
| --- | --- | --- | --- | --- |
|  | DMSO |  | TRiCi |  |
|  | positive strand | negative strand | positive strand | negative strand |
| gs 1 (VP1) | 1234.23 | 90.94 | 1874.40 | 26.46 |
| gs 2 (VP2) | 2055.77 | 128.91 | 2340.52 | 33.37 |
| gs 3 (VP3) | 414.28 | 117.13 | 936.93 | 31.11 |
| gs 4 (VP4) | 1021.22 | 36.40 | 1283.85 | 9.83 |
| gs 5 (NSP1) | 1167.30 | 261.97 | 275.04 | 58.00 |
| gs 6 (VP6) | 5392.04 | 251.83 | 5365.27 | 75.08 |
| gs 7 (NSP3) | 1195.27 | 183.01 | 1986.40 | 44.96 |
| gs 8 (NSP2) | 6346.81 | 116.86 | 6825.37 | 37.34 |
| gs 9 (VP7) | 7771.91 | 226.33 | 7526.26 | 77.71 |
| gs 10 (NSP4) | 5846.58 | 345.27 | 7324.93 | 108.73 |
| gs 11 (NSP5) | 7377.46 | 383.15 | 7548.64 | 131.79 |
| <sup>b)</sup> GAPDH | 7.47 | 0.00 | 4.33 | 0.00 |

<sup>a)</sup> MA104 infected with porcine strain OSU
<sup>b)</sup> Sequence covering GAPDH of *Macaca fascicularis*

### TRiC associates with NSP5 and VP2

As (-)ssRNA synthesis is impaired upon TRiC inhibition, we wondered if other viroplasm components are associated with TRiC in addition to NSP5. To assess this option, we took advantage of a well-described (66–69) *in vivo* protein-protein interaction assay based on the ability of the C-terminal region of reovirus protein µNS (amino acids 471-721) to form spontaneous cytosolic inclusions. In this particular assay, we used as ‘bait’ the red monomeric mCherry fused to the N-terminus of (471–721) µNS, herein mCh-µNS. The RV proteins (VP1, VP2, VP3, VP4, VP6, NSP2, and NSP5) were fused at the N-terminus of mCh-µNS **(Fig S7a)**. A specific antibody targeting the subunit CCT3 of TRiC was employed as ‘fish’ for the detection of TRiC localization. The co-localization, visualized by fluorescence microscopy, of the fish target protein with the RV protein-mCh-µNS cytosolic platform was considered as a positive protein-protein interaction. As expected **(Fig 6a)**, the ability of the CCT3 subunit to colocalize with the mCh-µNS resulted negative. However, CCT3 was clearly associated with VP1-mCh-µNS, VP2-mCh-µNS, and NSP5-mCh-µNS and only partially with NSP2-mCh-µNS and VP6-mCh-µNS. No association was observed for CCT3 with VP3-mCh-µNS, VP4-mCh-µNS, and VP6-mCh-µNS. Next, we immunoprecipitated TRiC from RV-infected MA104 cells at 6 hpi to confirm whether TRiC associates with the VP1, VP2, and NSP5. The resolved proteins showed that VP2 and NSP5 co-immunoprecipitated with TRiC when detected by immunoblotting using specific antibodies targeting these proteins (**Fig 6b**) but not VP1. Similarly, expression of either VP2 (**Fig 6c**) or NSP5 (**Fig 6d**) alone in BHK/T7 cells showed association of both proteins with TRiC by co-immunoprecipitation. However, TRiC did not co-immunoprecipitate V5-VP1 (**Fig 6e**). These results suggest that VP2 and NSP5, but not VP1, are the main RV proteins associated with TRiC.

**Figure 6.**
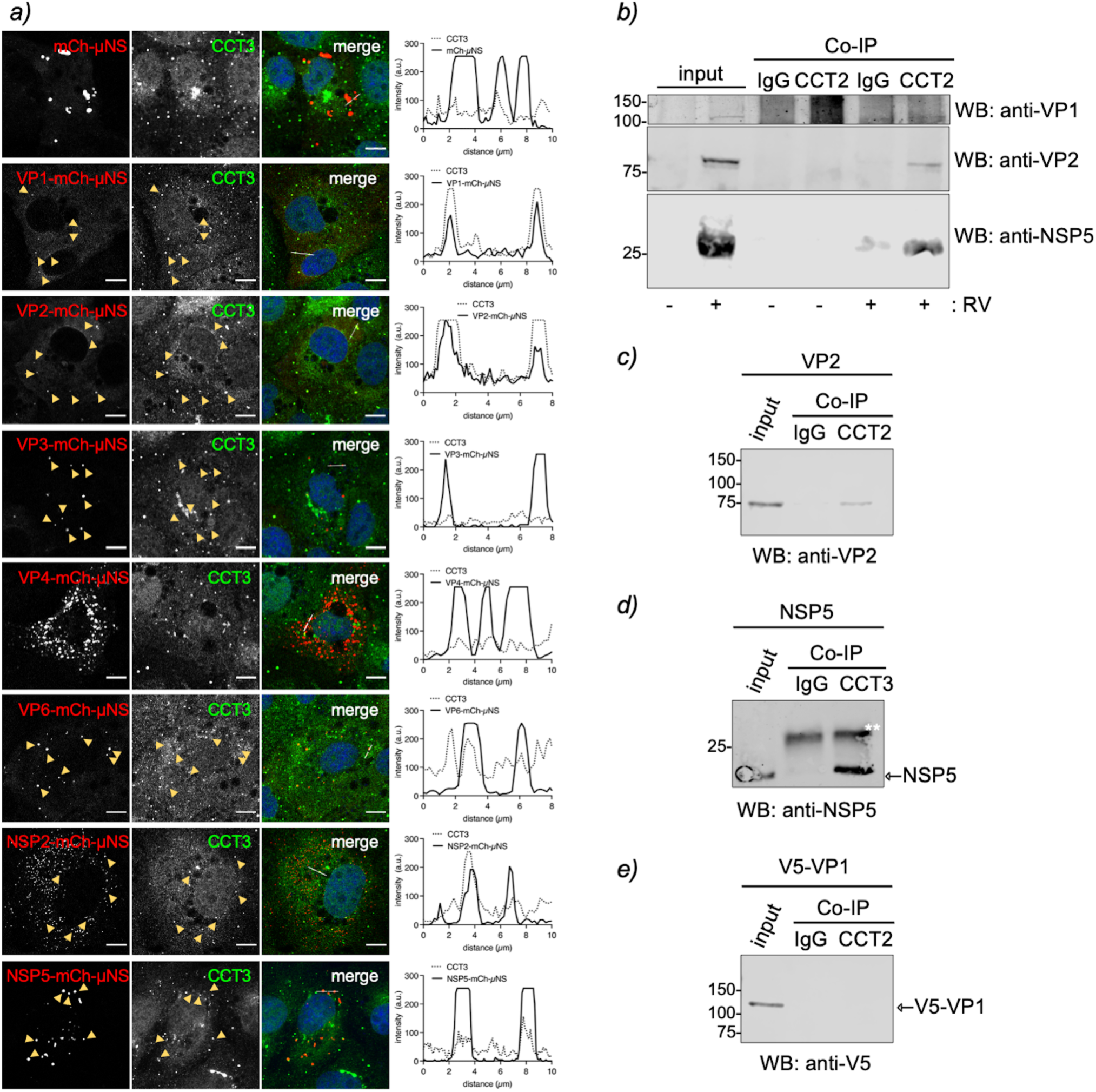
VP2 and NSP5 associate with TRiC. a) *In vivo* protein-protein interaction platform assay using as “bait” mCh-µNS fused at its N-terminal with the indicated RV-proteins. The “fish” interactor corresponds to the TRiC subunit, CCT3. At 24 hpt, the cells were fixed and immunostained for CCT3 (anti-CCT3, green). Yellow arrowheads point to mCh-µNS globular cytosolic platforms. Each plot corresponds to the intensity profile of the linear region of interest (LROI) indicated in the corresponding left image frame. **b)** Anti-CCT2 immunoprecipitated from OSU-infected MA104 cell extracts (6 hpi, MOI of 25 VFU per cell) and immunoblotted for detection of VP2 and NSP5. Anti-TRiC immunoprecipitated of BHK/T7 cell lysates expressing VP2 **(c)**, NSP5 **(d),** and V5-VP1 **(e)**. The membranes were incubated with the indicated antibodies. The input corresponds to 5% of crude cell extract. IgG corresponds to immunoprecipitation with isotype control antibody.** points to the light chain immunoglobulin.

Since VP2 and NSP5 are mainly found in viroplasms during the RV life cycle, we wondered if these two proteins have a role in accumulating TRiC in these inclusions. In this context, highly complex viroplasms can be studied through simplified structures termed viroplasm-like structures (VLS), which are based on the co-expression of the main viroplasm-building protein, NSP5 with either NSP2 or VP2 (12, 18). Therefore, we induced the formation of VP2 or NSP2 viroplasm-like structures (VLS) with other RV proteins present in viroplasms, such as VP6 and V5-VP1 (**Fig 7a and Fig S7b**). The diverse VLSs were monitored for the recruitment of the TRiC subunit CCT3 by confocal immunofluorescence, followed by quantification of the accumulation of the CCT3 signal in VLSs. All the combinations of built VLSs showed a similar in morphology based in homogenous NSP5 signal, a common marker for VLSs (**Fig 7b**). We noticed that CCT3 localization in VLSs was enhanced in VLSs containing VP2 but not in those containing with NSP2 (**Fig 7c**). Similarly, VLSs containing additional VP6 or V5-VP1 did not improve CCT3 accumulation in VLSs, suggesting no role of these proteins in the recruitment of TRiC. Notably, CCT3 showed a basal accumulation in VLSs composed of NSP5 and NSP2, consistent with the ability of NSP5 to associate with TRiC. Other TRiC subunits, such as CCT1 and CCT2, were also localized in VLSs containing NSP5 and either NSP2 or VP2 (**Fig 7d**). These results suggest that even if both VP2 and NSP5 associate with TRiC, VP2 is largely responsible for recruiting TRiC in the viroplasms.

**Figure 7.**
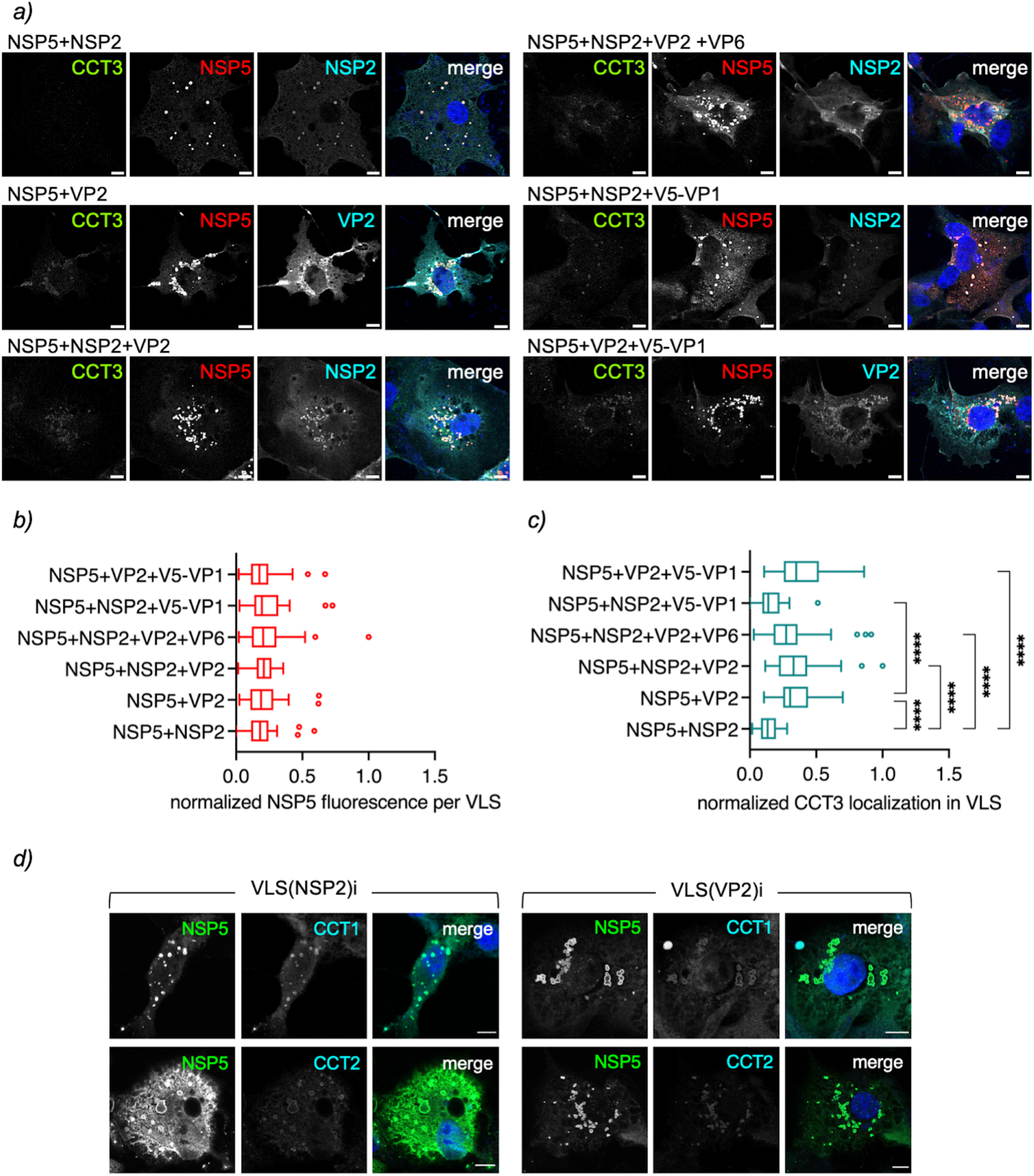
VP2 enhances the recruitment of TRiC components into VLSs. a) Immunofluorescence images of VLSs composed of the indicated RV proteins. At 16 hpt, the cells were fixed and immunostained to detect CCT3 (anti-CCT3, Alexa 488, green), VLS (anti-NSP5, Alexa 594, red), and VP2 (anti-VP2, Alexa 647, cyan) or NSP2 (anti-NSP2, Alexa 647, cyan). Nuclei were stained with DAPI (blue). Scale bar is 10 µm. Plots for quantifying NSP5 **(b)** and CCT3 **(c)** localization in VLSs composed of the indicated RV proteins. The data means were compared using the Tukey method where (*), p<0.05, and (****) p<0.0001. **d)** Immunofluorescence images of VLS composed either NSP5 with NSP2 (left panel) or NSP5 with VP2 (right panel). At 16 hpt, the cells were fixed and immunostained for the detection of VLS (anti-NSP5, Alexa 488, green) and the TRiC components CCT1 (top row; anti-CCT1, Alexa 647, cyan) or CCT2 (bottom row; anti-CCT2, Alexa 674, cyan). Nuclei were stained with DAPI (blue). Scale bar is 10 µm.

### VP2 folding process necessitates TRiC

We then wondered whether the folding of VP2 or any other RV viroplasm protein was TRiC-dependent. We designed an assay using TRICi and a proteasome inhibitor added to RV-infected cell cultures at different times post-infection **(Fig 8a)** and compared the level of expression of different virus proteins at 6 hpi following treatment with TRICi (at 1 hpi) **(Fig 8b)**, assuming that proteins that are TRiC-folding dependent would undergo proteasome degradation. Therefore, the protein expression will be underrepresented upon TRiC inhibition. Consequently, the addition of the proteasome inhibitor at 3.5 hpi should rescue their expression levels. Densitometry of the band intensities was compared to untreated cellular extracts (lane 1) in at least three independent experiments **(Fig 8c)**. As shown in **Fig 8b and c**, the only TRiC-folding dependent was VP2, which showed reduced expression in the presence of TRICi and was partially rescued by treatment with UBEI-41 (70), a ubiquitin-activating enzyme E1 inhibitor. To confirm this result **(Fig 8d)**, we expressed each RV protein alone and treated with TRICi and UBEI-41 at 3.5- and 7-hours post-transfection (hpt), respectively. The cell lysates were prepared at 22 hpt, and RV proteins were detected by immunoblotting using specific antibodies. Consistent with the previous result **(Fig 8e)**, the densitometric analysis revealed the dependence of VP2 on TRiC for folding. Meanwhile, the other RV proteins tested were not sensitive to TRiC/UBEI-41 treatment. Of note, in both assays, we found a decrease in the expression of some RV proteins following TRiC inhibition, which is not recovered after UBEI-41 treatment. This reduction in expression is associated with an indirect effect on its expression due to the inactivation of TRiC(71)(44). In addition, in the presence of TRiCi and YOD1(C160S) **(Fig 8f, lanes 3 and 4)**, a dominant negative mutant for deubiquitinase YOD1 (72), VP2 levels were even higher than in untreated samples **(Fig 8f, lane 4)**. In contrast, the expression of VP2 alone with TRICi resulted in a less intense band than in untreated samples **(Fig 8f, lane 2)**. Collectively, these results suggest that VP2 is the primary target for TRiC-assisted folding in the viroplasms.

**Figure 8.**
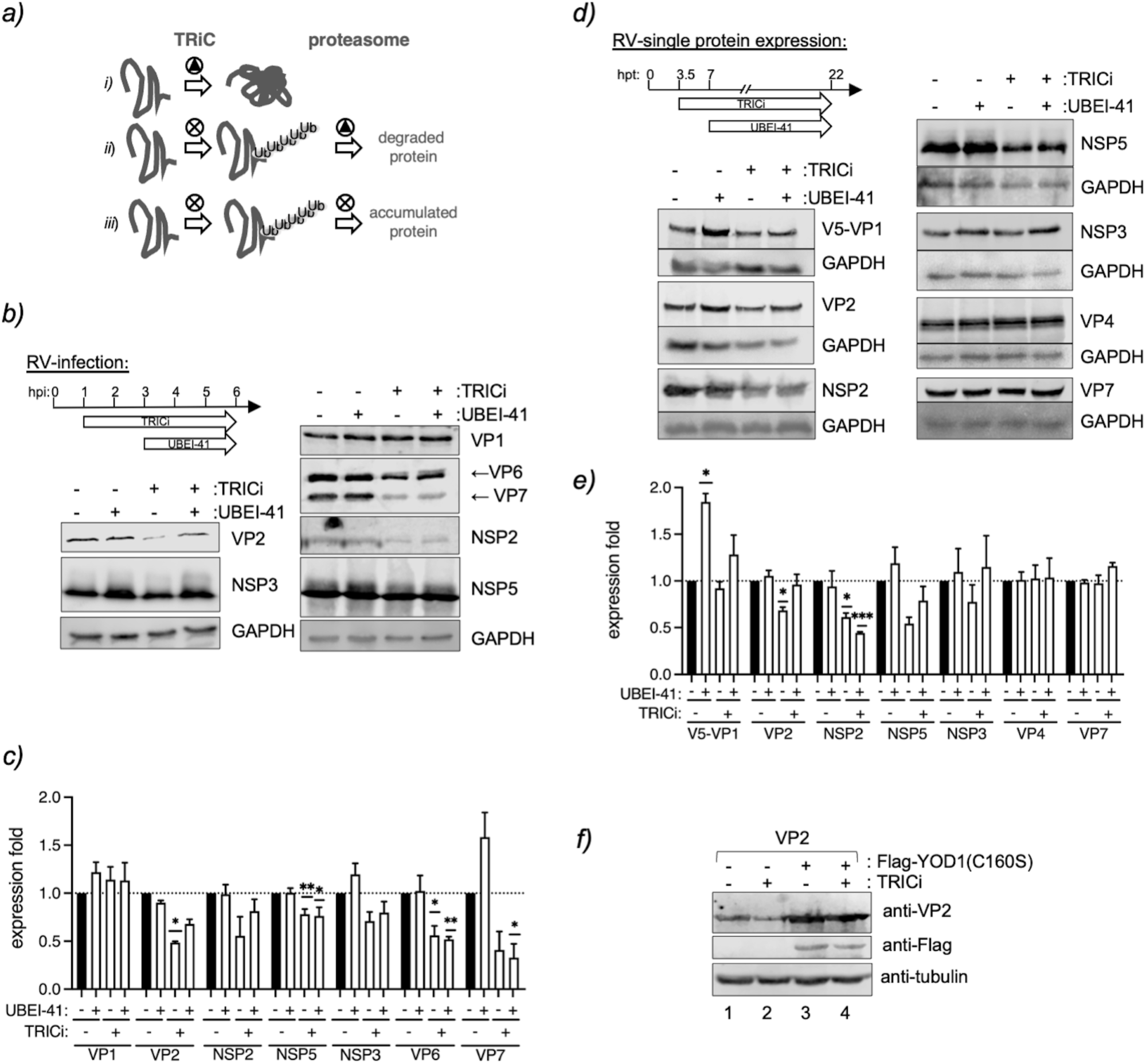
VP2 folding is TRIC-dependent. a) Scheme for the detection of TRiC folded RV proteins dependence. Cells expressing RV proteins (infection or transfection) are untreated or treated with a TRiC inhibitor (TRICi), followed by treatment with a proteasome inhibitor (UBEI-41). A TRiC folding dependent-RV protein in untreated conditions *(i)* is detected by immunoblot to an arbitrary expression value of 1. However, upon TRICi treatment *(ii),* its expression level decreases because unfolded protein degrades in the proteasome. If TRiC and proteasome are inhibited *(iii)*, the unfolded RV protein is not degraded and accumulates, being detected with higher levels than only the TRICi-treated sample. **b)** Immunoblotting showing the expression of RV protein at 6 hpi. RV-infected cells (MOI: 25 VFU/cell) were untreated or treated with 2.5 mM TRICi at 1 hpi, followed by treatment without or with 10 µM UBEI-41 at 3 hpi (top diagram). Next, the membranes were incubated with specific antibodies to detect the indicated RV proteins. **c)** Plot for the expression of the indicated RV proteins normalized to the corresponding untreated condition (black bar). The data correspond to at least three independent experiments analyzed with a one-sample t-test normalized to the corresponding untreated sample to a value of 1. (*), p<0.05 and (**), p<0.01. **d)** Immunoblotting showing RV proteins expressed in the absence of other virus proteins at 22 hpt. At 3.5 hpt, cells were untreated or treated with 2.5 mM TRICi followed by the addition, when indicated, of 10 µM UBEI-41 at 7 hpt (top diagram). Then, the membranes were incubated with specific RV antibodies. Except for V5-VP1, which was incubated with mouse mAb anti-V5. **e)** Plot for the expression of RV proteins in the absence of other virus proteins normalized to the corresponding untreated condition (black bars). The data correspond to at least three independent experiments analyzed with a one-sample t-test normalized to the corresponding untreated sample to a value of 1. (*), p<0.05 and (**), p<0.01. **f)** Immunoblotting of cellular extract co-expressing VP2 without or with Flag-YOD-1 (C160S), a dominant negative deubiquitinase, after 22 hpt. The cells were untreated or treated with 2.5 mM TRICi at 3 hpt. Then, the membranes were incubated with the indicated specific antibodies.

## Discussion

The role of TRiC in folding the mammalian orthoreovirus (MRV) σ3, another member of the Reoviridae family, has been recently demonstrated to be essential for the virus capsid assembly (55, 73). Using tandem mass spectrometry, we identified all TRiC subunits in viroplasms of RV and found that TRiC plays an essential role in the RV life cycle, specifically in the synthesis of the dsRNA genome segments. Moreover, we provide neat evidence that the inhibition of TRiC results in fewer viroplasms, lack of TLPs in the ER, and defective DLPs lacking encapsidated genome segments. Our result demonstrated that the inhibition of TRiC impairs (-)ssRNA RV synthesis but not (+)ssRNA. We also show that VP2 and NSP5 are associated with TRiC. The RdRp of RV, VP1, has a double task corresponding to transcription and replication. When transcribing, VP1 uses as a template (-)ssRNA for synthesizing (+)ssRNA, which is required for the translation of RV proteins but also as a template for replication of its genome segments. It is thought that transcription occurs in two waves (74, 75). The first transcription wave takes place immediately after internalization when transcriptionally active DLPs are released in the cytosol, permitting the translation of RV proteins required for halting host innate immunity, shutting off host translation, and building the viroplasms. The second wave of transcription occurs in the viroplasms, where the generated transcripts are released to the cytosol for translation by ribosomes (74, 76). The replication corresponding to the synthesis of dsRNA genome segments required for virus progeny occurs only in the viroplasms. Although it has been demonstrated by *in vitro* experiments the strict requirement of VP2 as a cofactor for VP1 to initiate the dsRNA synthesis (6, 77), no evidence has been provided in RV-infected cells. Moreover, high-resolution cryo-electron microscopy of purified RV particles deduced that the C-terminal plug of VP1 coordinates the replication process with VP2 (40, 41). The replication and packaging of each of the eleven genome segments in the new core-shell is a fine-tuned, not yet elucidated mechanism occurring in viroplasms. The current model (74) proposes a complex of VP1 and VP3 that associates in the viroplasms, in which VP1 interacts with the 3’ consensus sequence of each of the eleven (+) ssRNAs. In parallel, VP2 self-assembles forming decamers, allowing the concomitant formation of core shells and the recruitment of the VP1-VP3 complex. This process enables VP1 replication activity. In this context, our results provide new evidence on the RV genome replication and the packaging mechanism. Consistent with the model mentioned above, we demonstrate that VP2 improves the recruitment of TRiC in the viroplasms. Additionally, we demonstrate that TRiC can be found surrounding structures similar to DLPs. We provide evidence that TRiC mediates VP2 folding. Since VP2 directly associates with VP1 (78), and because the inhibition of TRiC impairs (-)ssRNA synthesis, we propose a model (**Fig 9, top**) in which TRiC coordinates the proper folding of VP2, which then associates with VP1 to initiate the genome replication process. In this model, TRiC would release properly folded VP2 able to associate with VP1 and to assemble into DLPs. The recruitment of TRiC in the viroplasms could result in a beneficial thermodynamic environment for VP2 to promote the initiation of the replication by VP1. The biochemical features of a substrate to be recognized by TRiC are poorly understood. Similar to MRV σ3, VP2 shares some biochemical features that could render it a suitable substrate. For example, the presence of conserved and complex beta-sheets in VP2 central and apical domains (79) suits with already described conditions for other beta-sheet-rich substrates like tubulin or actin (46). Alternatively, TRiC can stabilize VP2 higher-order structures favoring primed forms, allowing later association with VP1 and formation of the core shell. Thus, TRiC can retain a polypeptide in an almost native state until it binds to a protein interactor or a co-chaperone, such as Hsp70, to assist in the folding of higher-order structures (80, 81).

**Figure 9.**
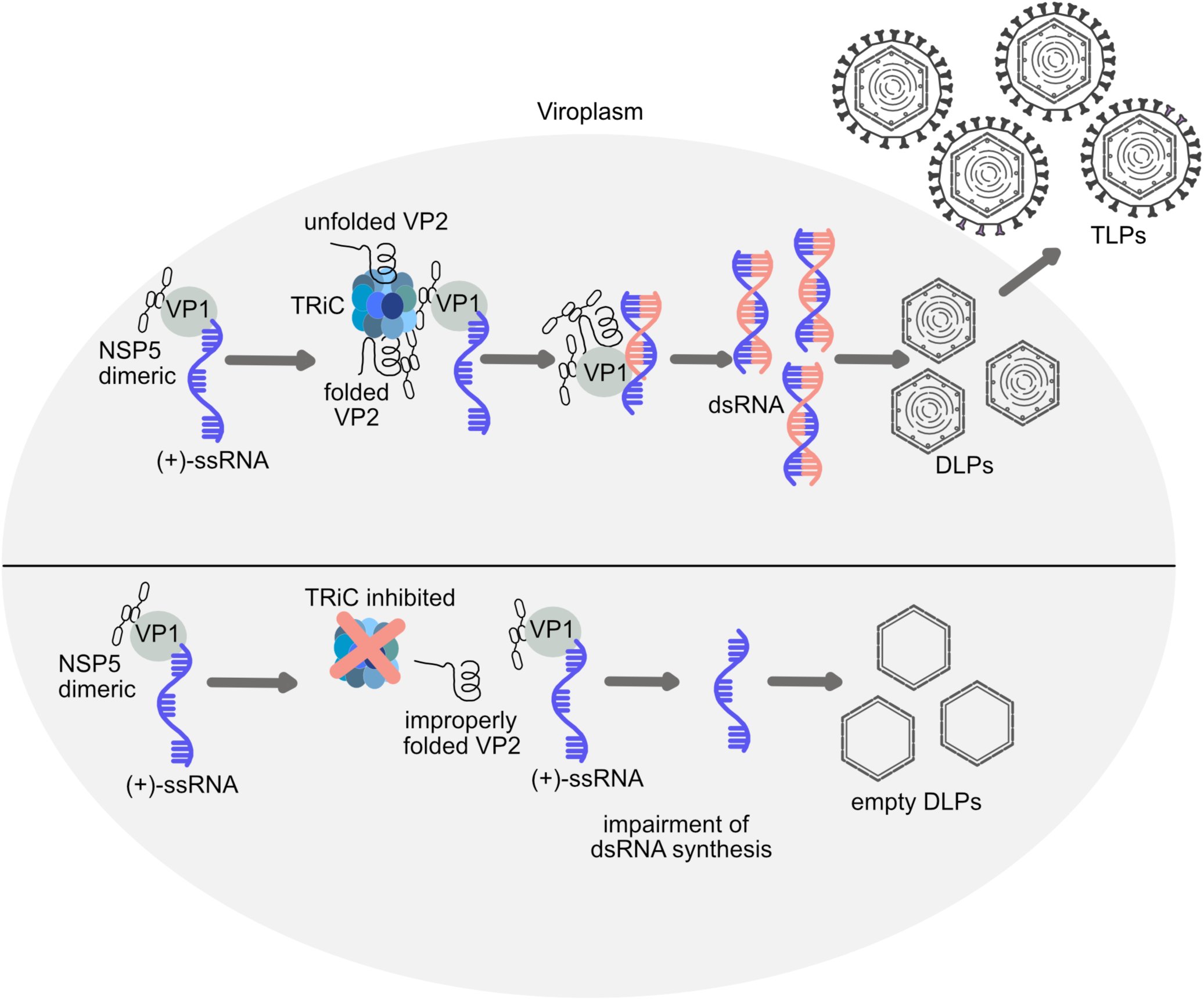
**Schematic model of the role of TRiC in RV replication**. As described in the top part scheme, the viroplasm is the site for the replication of the virus genome segments, specifically by synthesizing dsRNA using as template (+)ssRNA. In order to be accomplished this process, it is required that core shell protein VP2 assist the RdRp VP1 to shift from the synthesis of (+)ssRNA transcripts to the synthesis of dsRNA. This step is also obliged by NSP5. As a result of dsRNA genome segment synthesis, core particles are filled with eleven dsRNA genome segments to pursue the formation of DLP/TLPs. In contrast, when TRiC is inhibited (bottom part), VP2 is unfolded and, therefore, unable to assist VP1 switching to the synthesis of dsRNA genome segments. Due to the lack of genome replication, DLPs are empty of their genome segments, and TLPs are not generated.

The contribution of NSP5 also needs to be considered. Even if NSP5 folding shows to be independent of TRiC, we found NSP5 associates with TRiC and localizes in viroplasms surrounding DLP-like structures. As NSP5 has been previously demonstrated to be associated directly with VP1 (59) and bind RNA (82), as denoted in our model in **Fig. 9**, we cannot discard the possibility that NSP5 is involved in coordinating RV replication by mediation with TRiC. In this sense, an NSP5/KO recombinant RV is totally unable to replicate (7). Similarly, other RV proteins may depend on TRiC for folding. In fact, NSP3, which is not directly associated with viroplasms, was redistributed in the cytosol, forming small aggregates upon TRiC inhibition, an indicator for misfolded proteins.

This study provides evidence of the mechanism by which dsRNA genome segment replication is regulated and coordinated in the RV viroplasms.

## Materials And Methods

### Cells and viruses

MA104 cells (embryonic rhesus monkey kidney, ATCC^®^CRL-2378, RRID: CVCL_3845) were cultured in Dulbecco’s modified Eagle’s media (DMEM, Gibco^®^BRL) supplemented with 10% fetal calf serum (FCS)(AMIMED, BioConcept, Switzerland) and penicillin (100 U/ml)-streptomycin (100 µg/ml)(Gibco, Life Technologies). The MA104/NSP5-BioID2 cell line was generated using a lentiviral system. Briefly, HEK293T cells were maintained in DMEM (Life Technologies) supplemented with 10% FBS (Life Technologies) and 50 μg/ml gentamycin (Biochrom AG). Approximately 7×10^6^ HEK293T cells (human embryonic kidney, RRID: CVCL_0063) were seeded in a 10 cm^2^ tissue culture dish 24 hours before transfection. For each well, 2.4 μg of pMD2-VSV-G, 4 μg of pMDLg pRRE, 1.8 μg of pRSV-Rev, and 1.5 μg pAIP-NSP5-BioID2 were co-transfected with Lipofectamine 3000 (Sigma-Aldrich) according to the manufacturer’s instructions. After 48 h, the virus was collected and filtered with a 0.45-μm polyvinylidene fluoride filter. The virus stock was immediately used or stored at −80 °C. For lentiviral transduction, MA104 cells were transduced in six-well plates with 1 ml of lentiviral supernatant for 2 days. The positive cells were selected in 2 µg/ml puromycin. BHK-T_7/9_ (baby hamster kidney stably expressing T_7_ RNA polymerase) cells were kindly provided by Naoto Ito (Gifu University, Japan) (83) and cultured in Glasgow medium supplemented with 5% FCS, 10% tryptose phosphate broth (Sigma-Aldrich), 10% FCS, penicillin (100 U/mL)-streptomycin (100 μg/mL), 2% nonessential amino acids, and 1% glutamine.

Rotavirus porcine OSU strain (G5; P[9]), simian SA11 strain (G3; P6[1]), and rhesus RRV strain (G3; P5B[3]) were propagated in MA104 cells, as described previously (84). Virus titer was determined as described previously by Eichwald et al. (21) and expressed as viroplasm-forming units (VFU) per milliliter.

### Antibodies and reagents

Guinea pig anti-NSP5, guinea pig anti-NSP2, mouse anti-NSP2, guinea pig anti-VP2, guinea pig anti-VP1 and mouse scFV anti-NSP5 clone 1F2 were described previously (15, 16, 59, 85). Mouse monoclonal (mAb) anti-VP6 (clone 2F) was a gift from Dr. N. Mattion (CEVAN, Buenos Aires, Argentina). Mouse mAb anti-VP7 (clone 159), mouse anti-VP5 (clone 4G2), and mouse mAb anti-VP2 clone (3E8) were kindly provided by Harry B. Greenberg (Stanford University, CA, USA). Rabbit anti-NSP3 and mouse anti-VP1 were kindly provided by Susana López (UNAM, Mexico). Rabbit anti-NSP4 was kindly provided by Daniel Luque (ISCIII, Madrid, Spain). Rabbit anti-CCT1 was purchased at Invitrogen. Rabbit anti-CCT2 and rabbit polyclonal anti-CCT3 (A6547) were purchased at Abclonal. Goat anti-mouse conjugated to 6nm colloidal gold particles and goat anti-rabbit conjugated to 12 nm colloidal gold particles were purchased from Jackson ImmunoResearch Europe Ltd. Mouse anti-GAPDH (clone GAPDH-71.1, RRID: AB_1078991) was purchased from Sigma-Aldrich. Mouse monoclonal anti-V5 was purchased at Abcam. Rat anti-Histone H3 phosphorylated Ser 328 (clone HTA28) S28P-Alexa 647 (RRID: AB_397065) was purchased at Biolegend. Rabbit anti-cdc20 (RRID: AB_890558) was purchased at Bethyl Laboratories, Inc. Mouse monoclonal anti-dsRed2 (RRID: AB_1562589) was purchased at Santa Cruz Biotechnology, Inc.

Streptavidin-Dylight 488 was purchased at Invitrogen. Streptavidin-HRP was purchased at Sigma Aldrich.

MG132, Paclitaxel (taxol), and nocodazole were purchased from Sigma-Aldrich. UBEI-41 was purchased from NOVA biologicals. TRICi corresponds to 2-[(4-chloro-2λ4,1,3-benzothiadiazol-5-yl)oxy]acetic acid, (STK526585) was chemically synthesized at Vitas M Chemical (62).

### DNA plasmids

pMD2.G (Addgene plasmid #12259, RRID:Addgene_12259), pMDLg/pRRE (Addgene plasmid# 12251, RRID:Addgene_12251) and pRSV-Rev (Addgene plasmid#12253, RRID:Addgene_12253) were a gift from Didier Trono (86). pAIP(Addgene plasmid #74171, RRID:Addgene_74171) was a gift from Jeremy Luban (87).

pAIP-NSP5-BioID2 was prepared by ligation of NSP5-BioID2 fragment in pAIP within NotI and EcoRI restriction enzymes. NSP5-BiolD2 was synthesized as a gene block by Genscript. pcDNA-V5-VP1, pcDNA-VP1, pcDNA-VP2, pcDNA-NSP5, pcDNA-NSP3, pcDNA-VP4, pcDNA-VP7, and pcDNA-VP6 were previously described (16, 18, 59, 88).

pCI-VP1-mCherry-(471–721)µNS was completely synthesized at Genscript (Table S1). pCI-VP2-mCherry-(471–721)µNS, pCI-VP6-(471–721)mCherry were obtained by PCR amplification of pT7-VP6-SA11 using specific primers to insert *Xho*I and *EcoR*I, followed by ligation in-frame on those in pCI-mCherry-(471–721)µNS (68). pCI-VP3-mCherry-(471–721)µNS was obtained by PCR amplification of pT7-VP3 (SA11) using specific primers to insert *Xho*I and *Mlu*I, followed by ligation in-frame on those in pCI-mCherry-(471–721)µNS. pCI-VP4-mCherry-(471–721)µNS was obtained by PCR amplification of pT7-VP4 (SA11) using specific primers to insert *Nhe*I and *Mlu*I, followed by ligation in-frame on those in pCI-mCherry-(471–721)µNS. pCI-VP2-mCherry-(471–721)µNS, pCI-NSP5-mCherry-(471–721)µNS, pCI-NSP2-mCherry-(471–721)µNS were obtained by PCR amplification of pcDNA-VP2, pcDNA-NSP5, pcDNA-NSP2 using specific primers to insert *EcoR*I and *Mlu*I, followed by ligation in-frame on those sites in pCI-mCherry-(471–721)µNS. All the oligonucleotides were obtained from Microsynth AG, Switzerland, and listed in Table S1.

### Pull-down assay

MA-NSP5-BioID2 cells (5 x 10^6^) were infected with RV-SA11 (MOI of 5 VFU/ cell). For biotin labeling, cells were immediately washed after adsorption with PBS. Next, the media was replaced by DMEM supplemented with 10% FBS, 50 μg/ml gentamycin, and 200 µM biotin (BRAND) and incubated for 17 h at 37°C. Subsequently, the cells were gently washed once in PBS and then lysed in lysis buffer (50 mM Tris pH 8.0, 500 mM NaCl, 0.1 mM EDTA). The cell lysate was harvested in a 1.5 ml tube and centrifuged at 13,000 rpm for 1 minute at 4°C. Next, the supernatant was collected and incubated with 40 µl of Streptavidin Mag Sepharose (GE Healthcare Life Sciences) in the rotator wheel for 1 h at 4°C. The beads were subsequently washed three times with 500 µl of lysis buffer supplemented with 0.5 % SDS, three times with lysis buffer supplemented with 1 % NP-40, and three times with lysis buffer. The beads were then recovered and used for the downstream experiments.

For reverse pull-down assay, 2.4 x 10^6^ MA104/NSP5-BioID2 cells were RV-infected at MOI of 25 VFU/cell. At 1 hpi, media was replaced by media containing 100 µM biotin in DMEM serum-free. The cells were harvested at 6 hpi by detaching the cells with 5 mM EGTA in PBS and spun down for 2 min at 1500 rpm. The cellular pellet was resuspended in 2.5 ml of ice-cold ATP-depletion buffer (1 mM sodium azide, 2 mM 2-deoxyglucose, 5mM EDTA, 5 mM cycloheximide in PBS without calcium and magnesium) and then centrifuged at 300 x g for 5 min at 4 °C. The cell pellet was resuspended in 1ml lysis buffer B (50 mM HEPES-KOH pH 7.5, 100 mM KCl, 5 mM EDTA, 10% glycerol) supplemented with a protease inhibitor cocktail (Roche, Switzerland). Next, the cells were lysed by freeze and thaw three times, followed by dounce homogenization (70 strokes). The cell lysates were clarified by centrifugation at 17,000 x *g* for 10 min at 4°C. The cell lysate was dialyzed overnight at 4°C against PBS using a Slide-A-Lyser mini dialysis cassette (MWCO 3500, Thermo Fisher Scientific). For the input, 50 µl cell lysate was collected. The cell lysate was mixed with 1 vol of lysis buffer B supplemented with 2% BSA and 50 µl of Dynabeads M-270 Streptavidin (Thermo Fisher Scientific, cat 65305) and incubated at 4°C for 30 min with rotation. The beads were equilibrated in lysis buffer B supplemented with 2%BSA and incubated in wheel for 30 min at 4°C. The bead-bound complexes were washed three times with 500 µl ice-cold TRiC washes buffer (50 mM HEPES-KOH pH 7.5, 100 mM KCl, 5 mM EDTA, 10% glycerol, 0.05% NP-40), eluted with 20 µl SDS sample buffer, and resolved by SDS-PAGE.

### Mass spectrometry and proteomics

MA104/NSP5-BioID2 cells (1.5 ×10^8^) were infected with SA11 (MOI, 5), treated at 6 hpi with 100 µM biotin, and lysed 24 h post-transfection in TNN buffer [Tris–HCl 100 mM (pH[8), NaCl 250 mM, NP-40 0.5 % with cOmplete protease inhibitor cocktail (Roche)]. The pull-down of biotinylated proteins was performed using 100 µl StrAv Mag Sepharose™ (GE Healthcare) for each condition and incubated in a rotation wheel for 3h at 4°C. For mass spectrometry analysis, the washed biotin pull-downs were digested directly with trypsin (200 ng) in 20 μl of 20 mM triethyl ammonium bicarbonate pH 8.5 for 16 hours at room temperature. The supernatant was removed, the beads were washed once with 50 μl of 0.1 % formic acid, and the two fractions were pooled and concentrated using STAGE tips (89). The samples were resuspended in 10 μl of 0.1% formic acid and analyzed by LC-MS/MS using a NanoEASY LC (Thermo) coupled with an AMAZON ETD ion trap (Bruker Daltonics). The resulting spectra were searched against the human and rotavirus proteomes using the GPM (90). Results were filtered to remove all results with an e-value > 0.05. Statistical analysis and plots were performed using R (4.1.3). The figure was finalized in Adobe Illustrator 2022.

### Immunofluorescence

Samples were processed for immunofluorescence as described in detail by Vetter *et al.*, 2022 (29). Images were acquired at CLSM and processed with imageJ2 version 2.3.0/1.53q (Creative Commons license).

For native detection of antigens in RV TLPs, coverslips were pre-embedded with 50 µg/ml fibronectin in PBS, incubated for 30 min at room temperature (91), and rinsed once with PBS. The purified TLPs, corresponding to 2.5×10^6^ VFU, were diluted in 200 µl TBS (137 mM NaCl, 5 mM KCl, 7 mM Na_2_HPO_4_, 5.55 mM dextrose, 25 mM Tris pH7.4, 1 mM MgCl_2_ and 1 mM CaCl_2_) and incubated for 60 min at room temperature. The samples were fixed with 2% paraformaldehyde in PBS for 10 min at room temperature, washed three times with 0.1% triton X-100 in PBS, and blocked with 1% BSA in PBS for 20 min at room temperature. The coverslips were incubated with primary antibody diluted 1:100 and secondary antibody diluted 1:500. Samples were mounted in Prolong Gold (Thermo Fisher Scientific) and examined at LSCM Leica SP8 inverse.

### dsRNA genome segments extraction and electropherotype

MA104 cells seeded at a density of 5×10^5^ in a 60 mm tissue culture plate were reverse transfected with 10 nM of indicated siRNA and 10 µl Lipofectamine RNAiMAX Transfection Reagent (ThermoFisher Scientific) in a final volume of 5 ml as described above. At 48 hpt, cells were RV-infected (MOI, 25 VFU/cell). For this purpose, cell monolayers were washed once with phosphate-buffered saline (PBS) (137 mM NaCl, 2.7 mM KCl, 8 mM Na_2_HPO_4_, and 2 mM KH_2_PO_4_ pH 7.4) and adsorbed with 500 µl of diluted virus for 1 h at 4°C. Then, 3 ml of DMEM serum-free was added to the cells and incubated at 37°C. At 6 hpi, media was removed, and cells were harvested in 500 µl of RNA extraction buffer (0.5 % NP-40, 150 mM NaCl, 1.5 mM MgCl_2_, 10 mM Tris pH 7.4) in a 1.5 ml tube. Then, the dsRNA genome was extracted with 500 µl phenol-chloroform pH 4.6, followed by ethanol-sodium acetate precipitation (92). The pellet was resuspended in 20 µl of distilled water and mixed with 18 µl gel loading dye (New England Biolabs, Inc.). Samples were resolved in a 10% SDS-polyacrylamide gel, dsRNA genome segments stained with GelRed (Biotium), and visualized at Odyssey Fc Imager (LI-COR Biosciences).

The expression folds for each RV genome segment of purified particles correspond to the mean of three independent experiments of the ratio between the genome segment signal of interest and the signal of VP6 capsid for this sample obtained by immunoblotting.

### Immune and transmission electron microscopy

For immune electron microscopy, three confluent T-75 flasks of MA104 cells per sample were OSU-infected (MOI, 50 VFU/cell). For this purpose, the virus was adsorbed for 1h at 4°C and incubated in DMEM serum-free for 5.5 hpi at 37°C. Then, cells were incubated for 30 min with media containing 10 µM Taxol. At 6 hpi, cells were released with trypsin, harvested in complete medium, and spinned down at 1500 rpm for 2 min at room temperature. The cellular pellets were prepared according to the Tokuyasu method (93). Briefly, the cells were fixed with 4% formaldehyde at room temperature, washed several times with 0.1 M Na-phosphate, pH 7.4, and pelleted for 3 min at 13’000 rpm and 37°C in 12 % gelatin. The gelatin-embedded blocks immersed in 2.3 M sucrose were kept overnight at 4°C, mounted on ultramicrotome specimen holders (UC6, Leica Microsystems, Wetzlar, Germany), and frozen by plunging into liquid nitrogen. After trimming to a suitable block size and shape, 70 nm sections were cut at -120°C using a dry diamond knife (Diatome, Biel, Switzerland). Flat ribbons were picked up with a wire loop filled with a drop composed of 1 % methylcellulose, 1.15 M sucrose in 0.1 M Na-phosphate, pH 7.4, and transferred onto carbon-coated formvar films mounted on 100 hexagonal mesh/inch copper grids. For antigen retrieval, the sections were incubated for 1 h at 40°C with 0.1 Na-phosphate buffer pH 5.5, washed with 50 mM glycine, blocked with 1% BSA, and incubated with mouse monoclonal anti-NSP5 (clone 1F2) or mouse monoclonal anti-VP6 (clone 2F) at a dilution of 1:1 and rabbit anti-CCT3 (Abclonal), rabbit polyclonal anti-CCT2 (Abclonal) or rabbit monoclonal anti-CCT2 (Abcam) at a dilution of 1:5 at room temperatures for 90 minutes, washed several times with 0.1% BSA. Then, the samples were incubated for 45 min at room temperature with a goat anti-mouse antibody conjugated to 6 nm colloidal gold particles and a goat anti-rabbit antibody conjugated with 12 nm colloidal gold particles (Jackson ImmunoResearch Laboratories, Inc. West Grove, PA, USA). After incubation, the samples were washed with 0.1 M Na-phosphate pH 7.4 and distilled water and transferred to a mixture of 1.8 % methylcellulose and 0.4 % uranyl acetate. After 5 min incubation, the grids were looped out, and the solution excess was drained and air dried to obtain a remaining thin film on the grid.

For transmission electron microscopy images, MA104 cells were seeded at a density of 2×10^4^ cells per well in a 24-well tissue culture plate and then immediately added the sapphire discs. At 48 h post-seeding, the cells were RV-infected at MOI of 75 VFU/cell and at 1 hpi untreated or treated 1.25 or 2.5 mM TRICi. The cells were fixed at 6 hpi with 2.5 % glutaraldehyde in 100 mM Na/K phosphate buffer, pH 7.4 for 1 h at 4°C, and kept in 100 mM Na/K phosphate buffer overnight at 4°C. Afterward, samples were post-fixed in 1% osmium tetroxide in 100 mM Na/K phosphate buffer for 1 h at 4°C and dehydrated in a graded ethanol series starting at 70 %, followed by two changes in acetone, and embedded in Epon. Ultrathin sections (60 to 80 nm) were cut and stained with uranyl acetate and lead citrate.

For negative staining, the RV particles were adsorbed for 10 min on glow-discharged carbon-coated Parlodion films mounted on 300 mesh per inch copper grids (Electron Microscopy Science, Hatfield, PA, USA). Samples were washed once with distilled water and stained with saturated uranyl acetate (Fluka) for 1 min at RT. For calculation of the diameter of virus particles by negative staining, the area of each virus particle was calculated using Imaris software (version 2.1.0/1.53c; Creative Commons license) and then converted to the diameter as follows: d= 2 x √cA⁄nJ, where *A* is the area and *d* is the diameter of the particle, respectively.

All the samples were acquired in a transmission electron microscope (CM12; Philips, Eindhoven, The Netherlands) equipped with a charge-coupled-device (CCD) camera (Orius SC1000A 1; Gatan, Pleasanton, CA, USA) run with a Digital Micrograph software (Gatan) at an acceleration of 100 kV. The images were processed for publication using Image J (version 2.0.0-rc-69/2.52p)

Quantification of viroplasms.

Viroplasm size, number, and frequency were quantified previously(21, 28). The data were processed using Microsoft®Excel for Mac (version 16.61.1). Statistical analysis, unpaired parametric two-way ANOVA, and plots were performed using Prism 10 for macOS version 10.0.0 (131) (GraphPad Software, LLC).

### Quantification of protein signal in VLS

The fluorescence signal values in the VLSs were determined similarly, as described previously by Eichwald et al., 2020 (66). The intensity profile of a linear region of interest (LROI) was obtained using the ImageJ plot profile tool. The co-localization value was obtained by calculating the area below the curve of intensity profiles of both NSP5 VLS and other proteins. For this purpose, the Image J magic wand tool was used to provide the grey value intensity for each point, from which a protein signal percentage was obtained on the occupied NSP5-VLS signal area below the curve. The CCT3 normalized value from the area under the curve was obtained using the following formula:

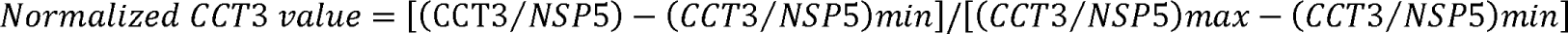

where CCT3 and NSP5 correspond to the value obtained of the area under the curve for each VLS. The minimal (min) and maximal (max) signal values for each condition were obtained from all the experimental tested samples.

### Immunoblotting

Cells seeded in twelve multi-well tissue culture plates at a density of 2×10^5^ cells per well were lysed directly by adding 25 µl of Laemmli sample buffer 4X (8 % SDS, 40 % glycerol, 200 mM Tris-HCl pH 6.8, 0.4 % bromophenol blue). The cell extracts were heated for 5 min at 95°C, sonicated for 5 sec at 14 Hz, and loaded in SDS-polyacrylamide gel. The proteins were migrated at 30 mA per gel and successively transferred to 0.45 µm Protan nitrocellulose membrane (Amersham). The membranes were blocked for 30 min in 5% milk-PBS and then incubated with primary and the corresponding secondary antibody conjugated to IRdye 680 or ICW780 (LI-COR). Samples were acquired at Odyssey Fc Imager (LI-COR Biosciences).

For expression of the proteins in RV purified particles, 1.2 x 10^6^ MA104 cells (three wells of a six-well multiwell plate) were infected with MOI of 12.5 VFU/cell. After 1 hpi, the cells were washed two times with PBS, and the media was exchanged by 1ml per well of media containing 2.5 mM TRICi in DMEM serum-free. At 8 hpi, media and cells were harvested and pooled in a 15 ml conical test tube. The samples were frozen in liquid nitrogen, thawed at 37°C in a water bath three times, and then clarified by centrifugation at 1500 rpm for 10 min at 4°C. The clarified supernatant was loaded over a 20% sucrose cushion in PBS using Ultra Clear tubes (13 x51 mm, cat n° 344057, Beckman Coulter) in a ratio of 3 to 4 of cushion and sample, respectively. The samples were ultracentrifuged for 27 Krpm for 2 h at 16°C using an AH-650 swinging bucket rotor (Thermo Scientific^TM^). Each pellet was resuspended in 50 µl of 50 mM Tris pH 7.5, 150 mM NaCl, and 0.1% Triton X-100. 5 µl of each sample was used to examine virus particles by transmitted electron microscopy, as described above. 22.5 µl of each sample was loaded in a 10% SDS-polyacrylamide gel and analyzed for immunoblot to detect RV proteins.

The expression value for each protein was obtained in a linear range protein analysis tool of Image Studio^TM^ Software (LI-COR Biosciences). The expression fold of RV proteins in cell extract was obtained by normalization to loading control (beta-actin). However, the expression fold of the RV proteins in purified particles was normalized with the VP6 capsid. The following formula was used:

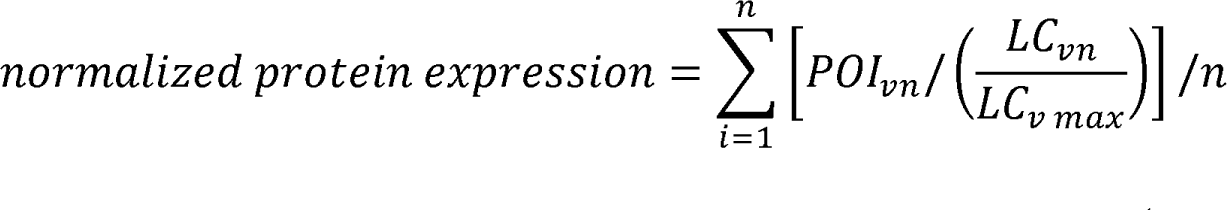

Where POI is the protein of interest, LC is the loading control (beta-actin or VP6 capsid), vn is the experimental value, and v max is the maximum value obtained for the loading control in each independent experiment. Similarly, for quantifying TRiC folding-dependent proteins after treatment with TRiC and proteasome inhibitors, the untreated samples were provided with a value of 1 for each independent experiment.

### Nanopore sequencing and data analysis

For extract preparation, MA104 cells were seeded at a density of 4×10^5^ cells/ well in six multiwell plates. The next day, the cells were washed twice with PBS and then infected with porcine strain OSU (MOI of 12.5 VFU/cell) diluted in 250 µl of DMEM serum-free. After 1 h of adsorption at 4°C, 750 µl DMEM serum-free were added to the well, and the cells were transferred to an incubator at 37°C. For experiments, including TRICi inhibitor, media was replaced at 1 hpi by adding 1.5 ml serum-free media containing either 2 % DMSO or 2.5 mM TRICi. At 6 hpi, the cells were lysed, and RNA was purified using an RNeasy mini kit (QIAGEN, Switzerland). Specifically, the genomic DNA was digested in-column using DNAse I as described by the manufacturer. RNA was eluted with 30 µl of nuclease-free water by incubating for 1 min before spinning down. Then, the column was re-eluted using the 30 µl elution. The RNA concentration and integrity were determined using 1 µl in 4150 TapeStation System (Agilent). The rest of the sample was immediately frozen at -80°C. For denaturation of dsRNA, the total purified RNA was heated for 5 min at 95°C and then immediately placed on ice. Then, the RNA sample was prepared as described at direct RNA sequencing protocol from Oxford Nanopore Technologies Limited, starting with an input of 1350 ng of RNA and a specific reverse transcription adapter (RTA). For this purpose, specific primers binding the 5’- and 3’-UTR of each RV genome segment and to GAPDH were synthesized at Microsynth AG (Switzerland) and described in Table S2. Oligo A was diluted to 2.8 µM TN buffer (10 mM Tris-HCl pH 7.5, 50 mM NaCl). Next, each oligo B, corresponding to RV 5’-and 3’-UTR of each genome segment, was diluted in TN buffer to 0.116 µM and pooled to reach a final concentration of 2.8 µM of total oligo B. Then, oligos were mixed 1:1 and annealed in a PCR machine (95° C for 2 min and then -1°C per 15 sec to 25°). This oligonucleotide was used as a replacement for RTA in the nanopore protocol. The samples were loaded on MinION Mk1B and started with the base calling in real-time using Guppy base calling software version (6.0.7+c7819bc) (Oxford Nanopore Technologies Limited). The data analysis was performed using Minimap2 software (94) followed by RStudio software version (2022.02.0+443) (RStudio: Integrated Development for R. RStudio, PBC, Boston, MA URL https://www.rstudio.com/).

The RNA sequence coverage from ONT is found in the SRA repository with the following accession reference: PRJNA900220 (https://www.ncbi.nlm.nih.gov/sra/PRJNA900220). The lead contact will share all data reported in this publication upon request.

### Co-immunoprecipitation of TRiC

For RV infection experiments, MA104 cells were seeded the day before at a density of 4 ×10^5^ cells/ well in 3 wells of tissue culture 6-multiwell plate and infected at MOI of 25 VFU/ cell. The RV-infected cells were harvested at 6 hpi. The day before transfection, BHK-T7 cells were seeded at a density of 4 ×10^5^ cells/ well in a tissue culture six-multiwell plate for single protein expression. The cells were transfected with 6 µg of pcDNA-VP2, pcDNA-NSP5, or pcDNA-V5-VP1 and 72 µl of Lipofectamine 2000 (Invitrogen) following the manufacturer instructions. The transfected cells were harvested at 24 hpt. The immunoprecipitation was adapted from the protocol described by Knowlton et al., 2018 (55). The cell monolayer was detached by adding 5mM EGTA in PBS and incubated for 10 min at 37°C. The cells were harvested in a 15 ml conical tube and spinned down for 5 min at 1500 rpm. The pellet was resuspended in 2.5 ml of ice-cold ATP-depletion buffer (1 mM sodium azide, 2 mM 2-deoxyglucose, 5 mM EDTA, 5 mM cyclohexamide in PBS without Ca^2+^ and Mg^2+^). Then, the cells were centrifuged at 300 *x* g for 5 min at 4 °C. The cell pellet was resuspended in 1ml lysis buffer B (50 mM HEPES-KOH pH 7.5, 100 mM KCl, 5 mM EDTA, 10% glycerol) supplemented with a protease inhibitor cocktail. The cells were lysed by freezing and thawing three times using liquid nitrogen and a 37°C water bath followed by dounce homogenization (70 strokes). The cell lysate was clarified by centrifugation at 17,000 x *g* at 4°C for 10 min and transferred to a new 1.5 ml tube. The input corresponded to 50 µl cell lysate. For immunoprecipitation, the cell lysate was split into two equal volumes and combined with 2 μg of rabbit CCT2 specific monoclonal antibody (Abcam, ab92746), 2 µg of rabbit anti-CCT3 (Ray Biotech, 144-06547-200) or 2 µg IgG isotype control antibody (Abcam, ab172730) and incubated at 4°C for 30 min with rotation. The cell lysates were then combined with 50 µl of Protein G Dynabeads (ThermoFisher, 10004D) equilibrated lysate buffer B and re-incubated at 4°C for 30 min with rotation. The bead-bound antibody-antigen complexes were washed four times with 500 µl ice-cold TRiC wash buffer (50 mM HEPES-KOH pH 7.5, 100 mM KCl, 5 mM EDTA, 10% glycerol, 0.05% NP-40), eluted with SDS sample buffer, and resolved by SDS-PAGE. The proteins were detected by immunoblotting, as described above.

### Statistical analysis

Statistical analyses were performed using Prism 9 (version 9.4.1, GraphPad Software) and RStudio software (version 2022.02.0+443, PBC, Boston, MA). The error bars and statistic tests are indicated for each corresponding experiment. *p* values of >0.05 were considered statistically significant (*, p<0.05; **, p<0.01; ***, p<0.001).

## Supporting information

Supplementary information: supplementary figure legends, description captions for tables S1and S2, supplementary materials and methods.

## Acknowledgments

This work was supported by the University of Zurich. A pre-doctoral ICGEB fellowship also supported this project for GP. The funders had no role in study design, data collection, interpretation, or the decision to submit the work for publication.

## Author contributions

Conceptualization, J.V., G.P., O.R.B., C.F. and C.E.; Methodology, J.V., G.P., K.T., M.M., O.R.B., C.F. and C.E.; Software, J.V. and K.T.; Validation, J.V., K.T. and C.E.; Formal analysis, J.V., K.T., M.M. and C.E.; Investigation, J.V., G.P., M.K., M.M., M.W., E.M.S. and C.E.; Resources, J.V., G.P., O.R.B., C.F. and C.E.; Data Curation, J.V., K.T. and C.E.; Writing-Original Draft, J.V. and C.E.; Writing-Review & Editing, J.V. and C.E.; Visualization; J.V. and C.E.; Supervision, O.R.B., C.F. and C.E.; Project Administration, C.E.; Funding Acquisition, O.R.B. and C.F.

