## Supplementary figures and images for "Recruitment of TRiC chaperonin in rotavirus viroplasms directly associates with virus replication"

### newFig S1.tiff

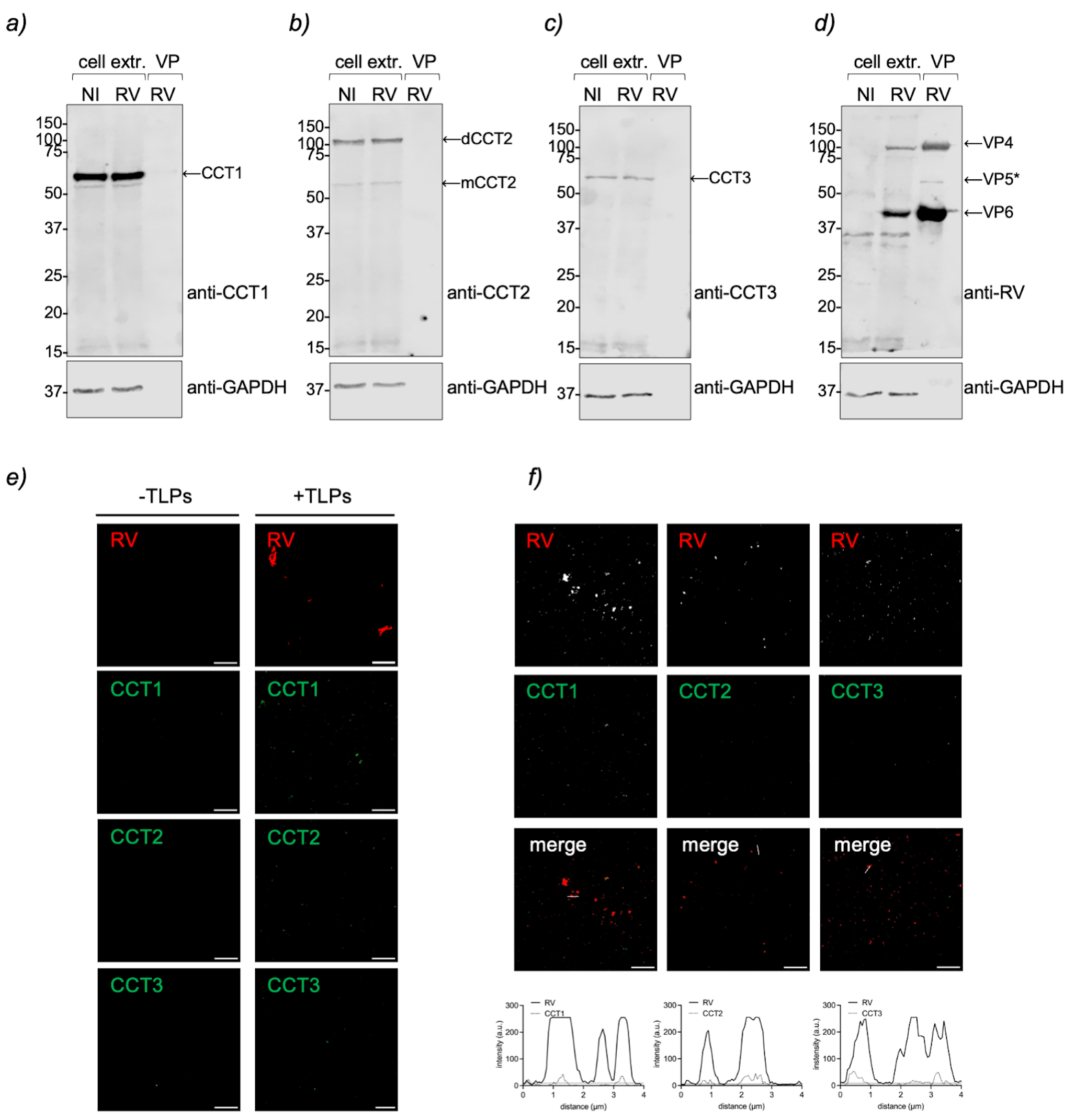

### newFig S2.tiff

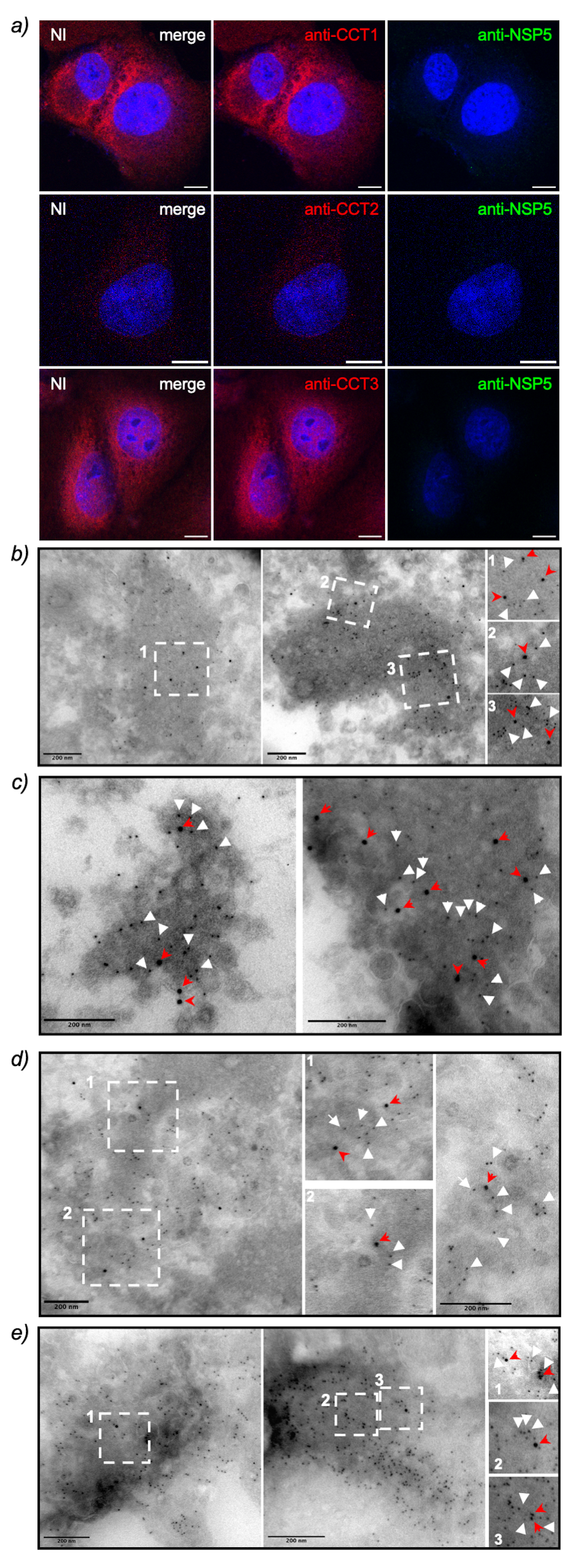

### newFig S3.tiff

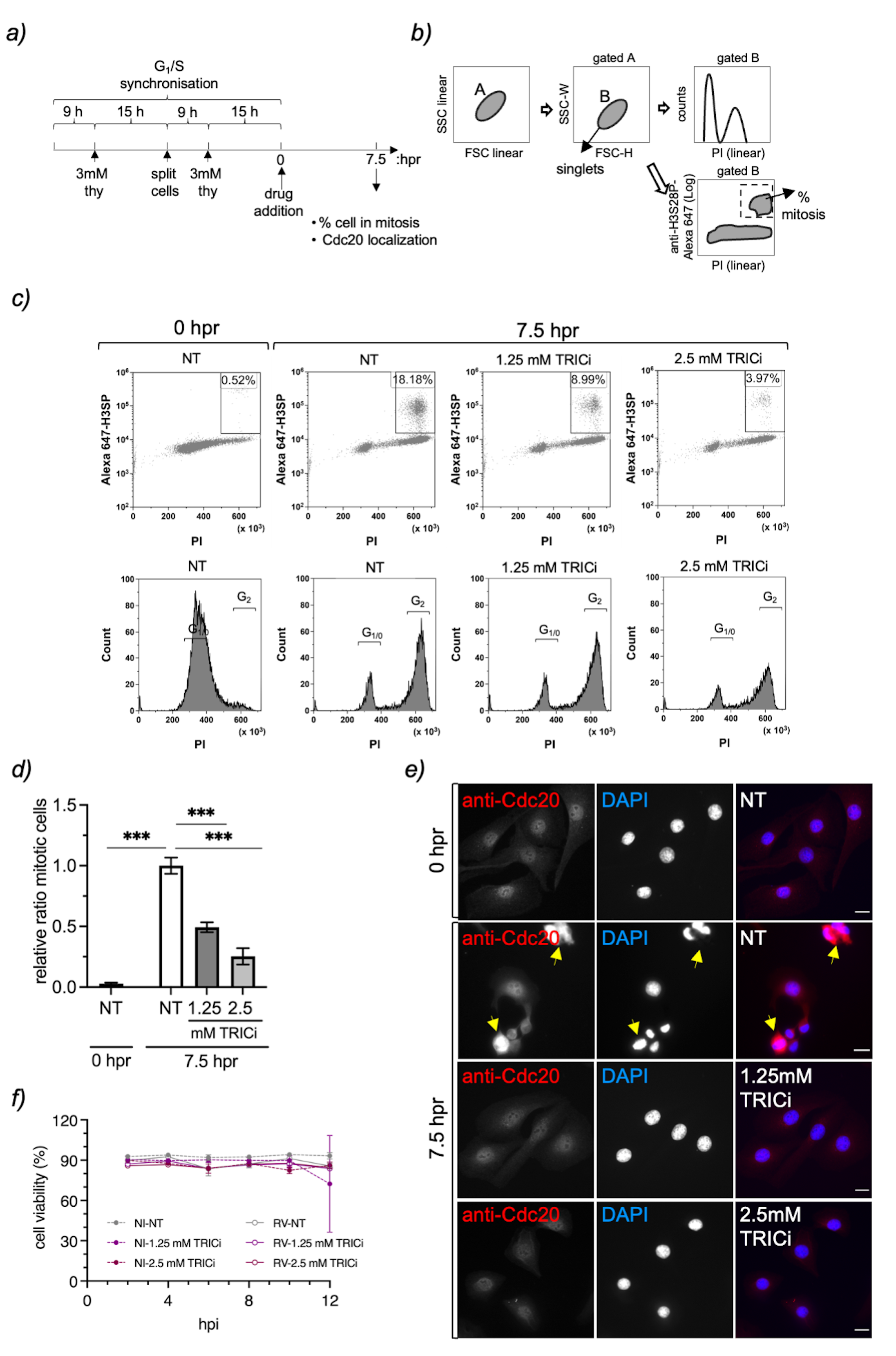

### newFig S4.tiff

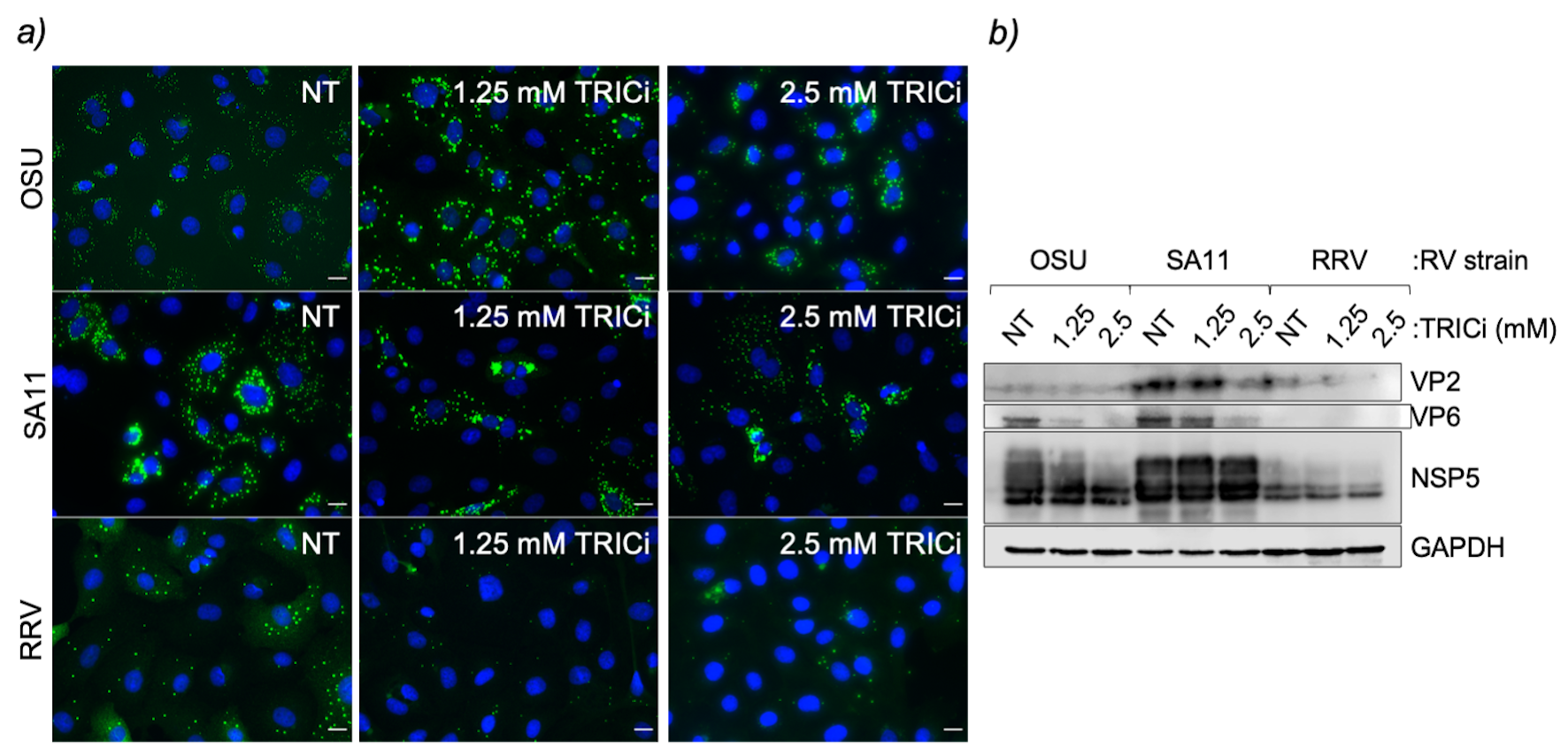

### newFig S5.tiff

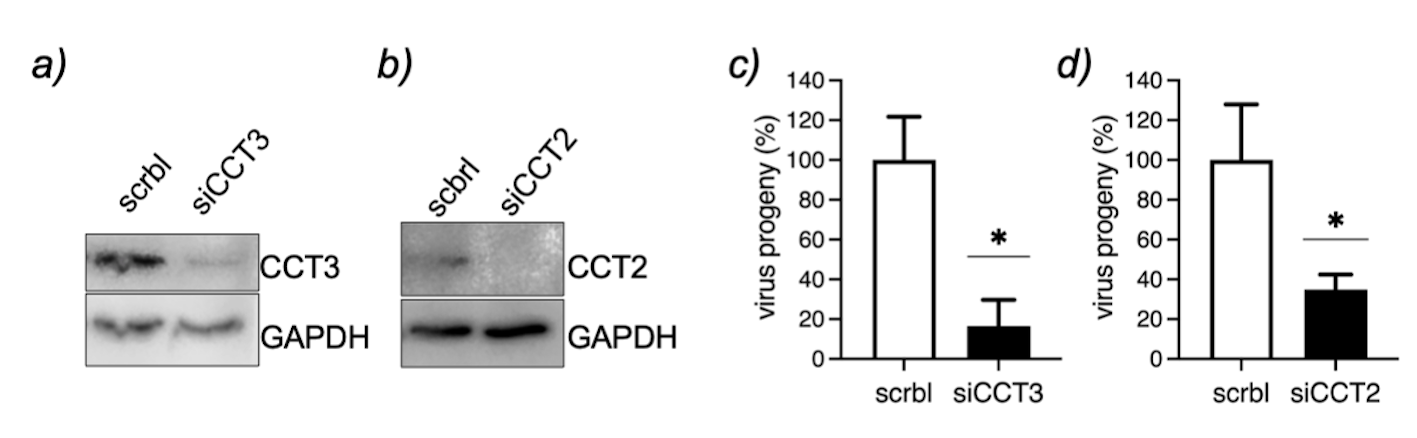

### newFig S6.tiff

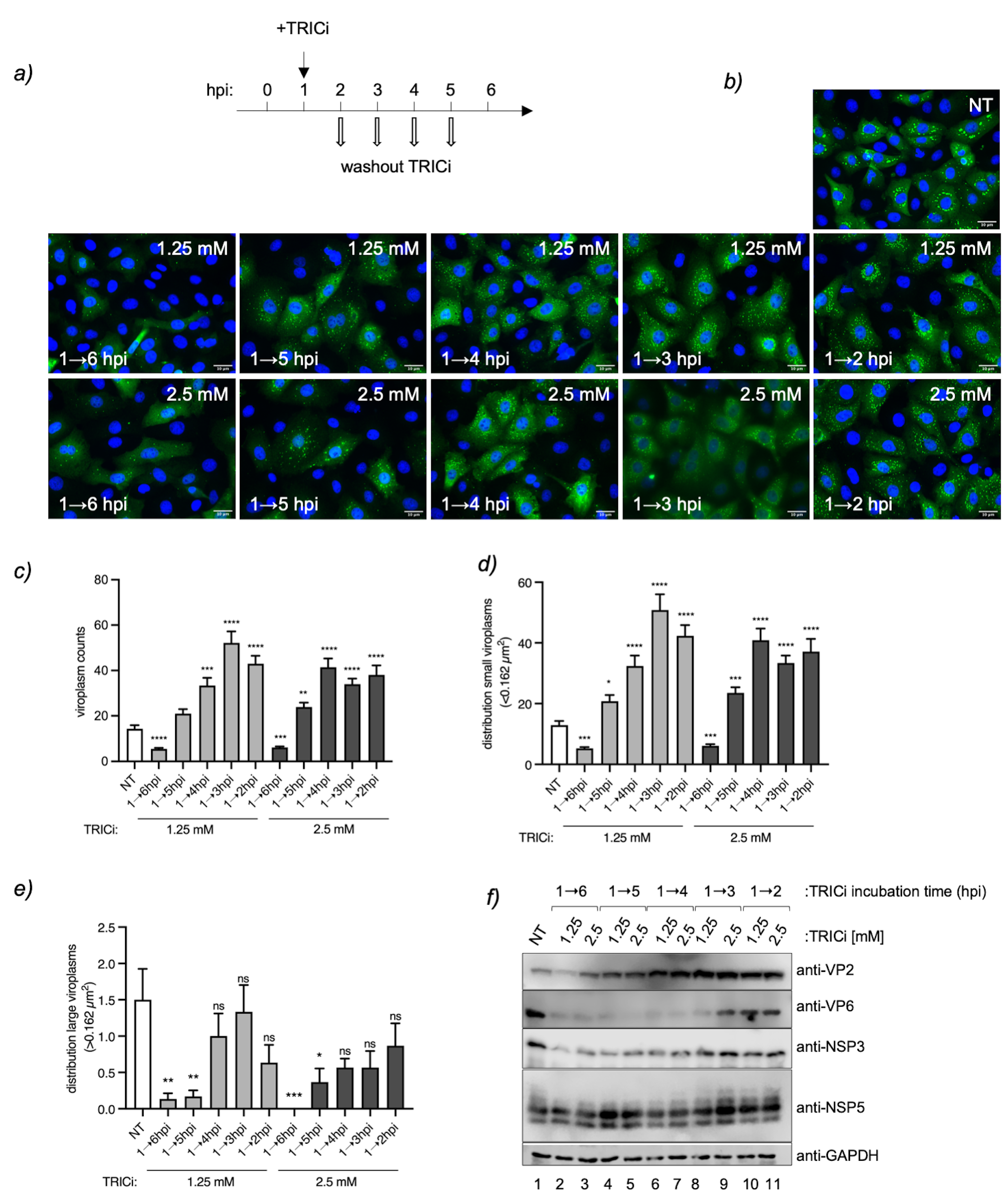

### newFig S7.tiff

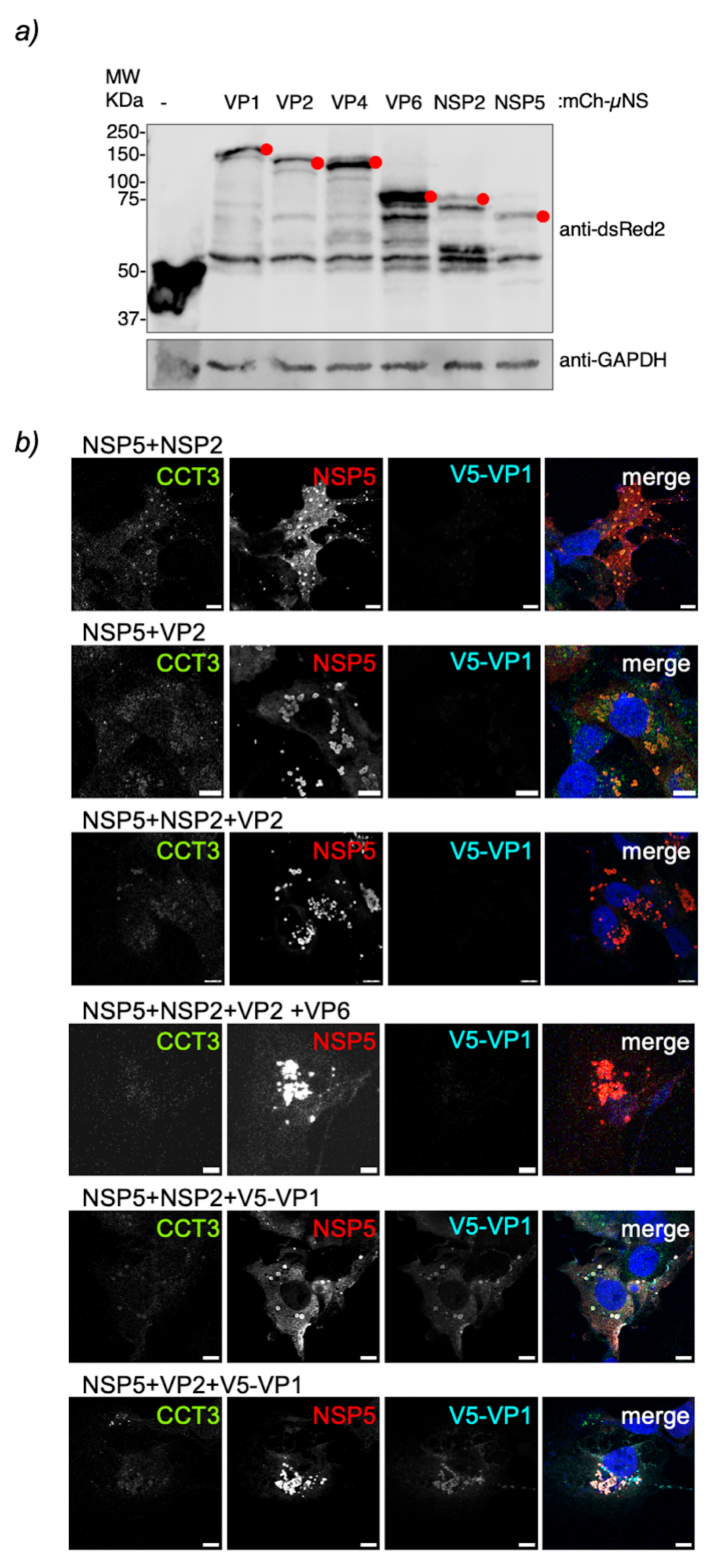
