## Supplementary information: supplementary figure legends, description captions for tables S1and S2, supplementary materials and methods. for "Recruitment of TRiC chaperonin in rotavirus viroplasms directly associates with virus replication": Supplementary Information.docx

**Supplementary Figure Legends**

**Figure S1. TRiC antibodies do not crossreact with RV antigens**. Immunoblot membranes of non-infected (NI) and RV-infected MA104 cell extracts and RV virus particles (VP) incubated with rabbit polyclonal anti-CCT1 **(a)**, rabbit polyclonal anti-CCT2 **(b)**, rabbit polyclonal anti-CCT3 **(c)** and guinea pig anti-RV **(d)** antibodies. The monomer and dimer of CCT2 are pointed with arrows. The RV proteins are indicated. GAPDH is used as a loading control. **e)** Immunofluorescence images of slides pre-embedded with fibronectin and coated without and with TLPs (2.5x10^6^ VFU) followed by immunostaining with guinea pig anti-RV (red), rabbit anti-CCT1 (green), rabbit anti-CCT2 (green) or rabbit anti-CCT3 (green). The scale bar is 10 µm. **f)** Immunofluorescence images of TLPs coated on slides pre-embedded with fibronectin followed by co-immunostaining with guinea pig anti-RV (red) and rabbit anti-CCT1, CCT2 or CCT3(green). A merged picture is shown in the right column. The scale bar is 10 µm. The plots correspond to the intensity profile of the linear region of interest (LROI) indicated in the bottom image frame.

**Figure S2. Distribution of CCT1, CCT2, and CCT3 in MA104 cells and the TRiC subunit CCT2 colocalizes in viroplasms surrounding DLPs. a)** Immunofluorescence images showing the distribution of TRiC subunits CCT1, CCT2, and CCT3 in non-infected MA104 cells. After fixation with methanol, the cells were co-immunostained with indicated specific antibodies for detecting TRiC subunits (Alexa 594, red) and anti-NSP5 (Alexa 488, green). Nuclei were stained with DAPI (blue). Scale bar is 10 µm. Immune-electron microscopy of viroplasm fixed at 6 hpi. The thin sections were co-immunostained with either anti-NSP5 conjugated to 6 nm gold **(b and c)** or anti-VP6 conjugated to 6 nm gold **(d and e)** followed by rabbit mAb anti-CCT2 **(b and d)** or rabbit polyclonal anti-CCT2 **(c and e)** both conjugated to 12 nm gold. The white dashed open boxes correspond to enlarged indicated images. Red arrowheads and white arrows point to the localization of CCT2 and NSP5 or VP6 surrounding DLPs. The scale bar is 200 nm.

**Figure S3. Characterization of TRICi in MA104 cells**. **a)** Schematic representation of MA104 cells synchronized in the G1/S phase using double thymidine blocking. Immediately after thymidine release, the cells were untreated or treated with TRICi at 1.25 mM and 2.5 mM. At 7.5 h post-release, cells were harvested and examined for the percentage of cells in mitosis using an anti-histone H3 phosphorylated serine 328 (Alexa 647-H3SP) and expression of Cdc20. **b)** Flow cytometry diagram to characterize the percentage of cells in mitosis. **c)** Representative scatter plots for mitosis detection (upper panel) and histogram plots for DNA content (lower panel) of MA104 cells untreated or treated with TRICi at the indicated concentrations. The percentage of cells in mitosis is indicated in the top right corner of each scatter plot. In addition, the G_1/0_ and G_2_ cell cycle phases are indicated. **d)** Plot for the relative ratio of mitotic cells after treatment with TRICi. The data correspond to the mean ± SD of three independent experiments. Two-way ANOVA, (***), p<0.0001. **e)** Immunofluorescence images for the expression of Cdc20 (anti-Cdc20, red) of thymidine synchronized MA104 cells followed by treatment without or with TRICi at the indicated concentrations immediately after release. Nuclei were stained with DAPI (blue). The yellow arrows point to Cdc20-positive cells. The scale bar is 10 µm. **f)** Plot for cell viability of non-infected and RV-infected MA104 cells untreated or treated with TRICi at 1.25 or 2.5 mM. The chemical compound was added at 1 hpi, and samples were collected up to 12 hpi. The cell viability was determined by the release of lactate dehydrogenase. The data represent the mean ± SD of three independent experiments. Positive control corresponds to cell lysed in 0.1 % Triton X-100 buffer.

**Figure S4. TRIC inhibition hampers viroplasm formation of RV strains OSU, SA11, and RRV.** OSU-, SA11-, or RRV-infected (MOI, 25 VFU/ml) MA104 cells untreated (NT) and treated with 1.25 mM or 2.5 mM TRICi added at 1 hpi. **a)** Immunofluorescence images of cells fixed at 6 hpi and immunostained to detect viroplasms (anti-NSP5, green). Nuclei were stained with DAPI (blue). The scale bar is 10 µm. **b)** Immunoblotting of cell extract harvested at 6 hpi. The membrane was incubated with specific antibodies to detect the indicated proteins. GAPDH is used as a loading control.

**Figure S5. Silencing CCT3 and CCT2 TRiC subunits decreases virus progeny and alters viroplasm morphology.** Immunoblotting of cellular extract silenced with siCCT3 **(a)** and siCCT2 **(b)**. Scrambled siRNA (scrbl) is included. GAPDH was used as the loading control. Plot for virus progeny of OSU-infected MA104 cells silenced with scrambled siRNA, siCCT3 **(c),** or siCCT2 **(d)**. The data represent the mean ± SD of three independent experiments. Student's t-test; (*), p<0.05.

**Figure S6. TRICi effect is reversible over viroplasm formation**. **a)** Schematic timeline for the addition and washout of TRICi in RV-infected cells. At 6 hpi, all the samples were analyzed to detect viroplasms and the expression of diverse RV proteins. **b)** Immunofluorescence images of RV-infected cells (MOI, 25 VFU/cell) treated with TRICi at indicated periods. At 6 hpi, cells were fixed and immunostained to detect viroplasms (anti-NSP5, green). Nuclei were stained with DAPI (blue). The scale bar is 10 µm. **c)** Plot for quantifying the number of viroplasms after recovery from TRICi as described in **(a)**. Plots for the distribution of small (< 0.162 µm^2^) **(d)** and large (> 0.162 µm^2^) **(e)** viroplasms after recovery from TRICi. Data represent the mean ± SD. n> 50 cells. The data represent the mean ± SD. n>50 cells per point, Welch's two-way ANOVA compared to untreated condition where (**), p<0.001; (***), p<0.0001 and (****), p<0.00001. **f)** Immunoblotting of RV-infected cell extracts after recovery at the indicated treatment periods with TRICi. The membranes were incubated with the indicated specific antibodies. GAPDH was used as a loading control.

**Figure S7. RV proteins associated with TRiC.** **a)** Immunoblot of cellular extracts expressing RV proteins fused to mCh-µNS. The membrane was incubated with the indicated antibodies. GAPDH was used as a loading control. The red dots point to the predicted molecular weight for each fusion protein. **b)** Immunofluorescence images of VLSs composed of NSP5 with the indicated RV proteins. At 16 hpt, the cells were immunostained for detection of VLS (anti-NSP5, red), TRiC subunit CCT3 (anti-CCT3, green), and V5-VP1 (anti-V5, cyan). The nuclei were stained with DAPI (blue). A merged image is shown in the right column. The scale bar is 10 µm.

**Supplementary Tables**

**Table S1**. Synthetic primers used in this study

**Table S2.** Reverse transcriptase adaptors used for Oxford Nanopore Technology direct RNA sequencing of RV positive and negative single-stranded RNAs

**Supplementary Materials and Methods**

**siRNA reverse transfection and infection.** siRNA reverse transfections were performed using Lipofectamine RNAi MAX Transfection Reagent following manufacturer instructions. For transfection in a 24-multiwell tissue culture plate, 1.2 µl siRNA 5µM were diluted to 100 µl with Opti-MEM™ (Gibco™, ThermoFisher) plus 1 µl of transfection reagent were added to one well and incubated for 20 min at room temperature. Then, 2x10^4^ MA104 cells diluted in 500 µl DMEM supplemented with 10%FCS were added on the top of the transfection. The siRNA final concentration reached is 10 nM. At 48 hpt, cells were RV-infected at an MOI of 25 VFU per cell, as described previously (1). The scramble siRNA corresponds to Control siRNA-A (sc-37007; Santa Cruz Biotechnology, Inc). The siCCT3 corresponds to the following equimolar mix of RNA sequences: CCT3_5 siRNA: 5’-CAGACTGACATTGAGATTACA-3'; CCT3_9 siRNA: 5’-AGCGGCCAAGTCCA TGATCGA-3'; CCT3_8 siRNA: 5’-ATCCACGTATGCGGCGCTATA-3'; CCT3_7: 5'- CTTGCGTGGAGTCATGATTAA-3' were purchased at Sigma-Aldrich.

**Cell synchronization and flow cytometry.** MA104 cells were synchronized by double thymidine blocking, as described previously by Gluck et al., 2017 (2). Cells were released by washing twice with PBS and adding 3 ml of 10% FCS-DMEM without or with TRICi at the indicated concentrations. At 7.5 h post-release, cells were harvested by washing twice with PBS, detached with 0.5 ml of 0.5 % Trypsin-EDTA (ThermoFisher), collected in 15 ml tubes with 3 ml cDMEM, and centrifuged for 2 min at 1500 rpm. The cell pellet was resuspended in 1 ml PBS and mixed gently with 2.5 ml ethanol 100%. The samples were stored at -20ºC overnight. Then, samples were centrifuged for 5 min at 1500 rpm at 4ºC, and the pellet was washed once with 1 ml of PBS, followed by a second wash with 1 ml of staining buffer (1% Fetal bovine serum in PBS). The cell pellet was resuspended in 100 µl of Alexa 647-anti-histone H3-Phosphorylated (Ser38) antibody (clone HTA28) (BioLegend) diluted 1:20 in staining buffer and incubated for 20 min at room temperature. Cells were centrifuged for 5 min at 1500 rpm and 4ºC, and the pellet was resuspended in 500 µl of propidium iodide solution (0.05 % Triton X-100, 0.1 mg/ml RNAse A, 50 µg/ml PI in PBS), incubated for 40 min at 37ºC in the dark and filtered using a cell strainer snap-cap tube (BD Falcon™). Samples were immediately acquired using a CytoFLEX S flow cytometer (Beckman Coulter). Thus, 25'000 events were acquired exciting at both 488 nm with a blue laser and filter band of 610/20 and 638 nm with a red laser and filter band 660/10. Data were analyzed and processed using Kaluza Analysis Software (Beckman Coulter). For the detection of Cdc-20 by immunofluorescence, a coverslip was included in the second blocking of thymidine. At the indicated time post-release, the cells in slides were fixed in 2% paraformaldehyde for 10 min at room temperature, permeabilized with 0.1% Triton X-100 in PBS for 5 min at room temperature and blocked with 1% BSA-PBS for 20 min at room temperature. Expression of Cdc20 was detected by immunostaining with a rabbit anti-Cdc20 followed by a secondary antibody anti-rabbit conjugated to Alexa 594. Nuclei were stained with 0.01 mg/ml DAPI in PBS. Coverslips were mounted in ProLong™ Gold Antifade Mountant (ThermoFisher). Images were acquired at CSLM SP8 inverse (Leica) using a 63X HCPL APO CS2 lens and analyzed using ImageJ2 version: 2.3.0/1.53q.
