## Supplementary material for "Recruitment of TRiC chaperonin in rotavirus viroplasms directly associates with virus replication": Table S1: Table S1.pdf

**Table S1.** Synthetic primers used in this study

| Amplified fragment | Oligonucleotide sequences |
|---|---|
| *Xho*I-VP6(SA11)-*EcoR*I | fwd: 5′-gatcgaattctttaatgagcatgcttctaat-3′<br>rev: 5′-gatcctcgagatggatgtcctatactctttg-3' |
| *Xho*I-VP3(SA11)-*Mlu*I | fwd: 5′-gatcctcgagatgaaagtactagctttaaga-3′<br>rev: 5′-gatcacgcgtctcaaccatatcaaacactgt-3' |
| *Nhe*I-VP4(SA11)-*Mlu*I | fwd: 5′-ctataggctagcgccaccatggcttcgctcatttatag-3′<br>rev: 5′-ggtaccacgcgtcaacctgcattgcataatcag-3′ |
| *EcoR*I-NSP5(OSU)-*Mlu*I | fwd: 5'gatcgaattcatgtctctcagcattgacgta-3'<br>rev: 5′-gatcacgcgtcaaatcttcgatcaattgcat-3′ |
| *EcoR*I-NSP2(SA11)-*Mlu*I | fwd: 5′-gatcgaattcatggctgagctagcttgcttt-3'<br>rev: 5′-gatcacgcgtaacgccaacttgagaaacttc-3' |
| *EcoR*I-VP2(SA11)-*Mlu*I | fwd: 5′-gatcgaattcatggcgtatcgaaaacgtgga-3'<br>rev: 5′-gatcacgcgtcagttcgttcatgatgcgcat-3' |

a) Inserted restriction enzymes are underlined.
