## Supplementary material for "Recruitment of TRiC chaperonin in rotavirus viroplasms directly associates with virus replication": Table S2: Table S2.pdf

**Table S2**: Reverse transcriptase adaptors used for Oxford Nanopore Technology direct RNA sequencing of RV positive and negative single stranded RNAs

| **name** | **oligonucleotide annealing positive strand** | **oligonucleotide annealing negative strand** |
|---|---|---|
| gs 1_(VP1) | 5′-*gaggcgagcggtcaattttcctaagagcaagaagaagcc***ggtcacatct**-3′ | 5′-*gaggcgagcggtcaattttcctaagagcaagaagaagcc***ggctattaaa**-3′ |
| gs2_(VP2) | 5′-*gaggcgagcggtcaattttcctaagagcaagaagaagcc***ggtcatatct**-3′ | 5′-*gaggcgagcggtcaattttcctaagagcaagaagaagcc***ggctattaaa**-3′ |
| gs3_(VP3) | 5′-*gaggcgagcggtcaattttcctaagagcaagaagaagcc***ggtcacatcg**-3′ | 5′-*gaggcgagcggtcaattttcctaagagcaagaagaagcc***ggctattaaa**-3′ |
| gs4_(VP4) | 5′-*gaggcgagcggtcaattttcctaagagcaagaagaagcc***ggtcacaacc**-3′ | 5′-*gaggcgagcggtcaattttcctaagagcaagaagaagcc***ggctataaaa**-3′ |
| gs5_(NSP1) | 5′-*gaggcgagcggtcaattttcctaagagcaagaagaagcc***ggtcacattt**-3′ | 5′-*gaggcgagcggtcaattttcctaagagcaagaagaagcc***ggcttttttt**-3′ |
| gs6_(VP6) | 5′-*gaggcgagcggtcaattttcctaagagcaagaagaagcc***ggtcacatcc**-3′ | 5′-*gaggcgagcggtcaattttcctaagagcaagaagaagcc***ggctttaaaa**-3′ |
| gs7_(NSP3) | 5′-*gaggcgagcggtcaattttcctaagagcaagaagaagcc***ggtcacataa**-3′ | 5′-*gaggcgagcggtcaattttcctaagagcaagaagaagcc***ggcttttaat**-3′ |
| gs8_(NSP2) | 5′-*gaggcgagcggtcaattttcctaagagcaagaagaagcc***ggtcacataa**-3′ | 5′-*gaggcgagcggtcaattttcctaagagcaagaagaagcc***ggcttttaaa**-3′ |
| gs9_(VP7) | 5′-*gaggcgagcggtcaattttcctaagagcaagaagaagcc***ggtcacatca**-3′ | 5′-*gaggcgagcggtcaattttcctaagagcaagaagaagcc***ggctttaaaa**-3′ |
| gs10_(NSP4) | 5′-*gaggcgagcggtcaattttcctaagagcaagaagaagcc***ggtcacacta**-3′ | 5′-*gaggcgagcggtcaattttcctaagagcaagaagaagcc***ggcttttaaa**-3′ |
| gs11_(NSP5) | 5′-*gaggcgagcggtcaattttcctaagagcaagaagaagcc***ggtcacaaaa**-3′ | 5′-*gaggcgagcggtcaattttcctaagagcaagaagaagcc***gcgctacagt** |
| GAPDH_MA104 | 5′-*gaggcgagcggtcaattttcctaagagcaagaagaagcc***ggggatttta**-3′ | 5′-*gaggcgagcggtcaattttcctaagagcaagaagaagcc***ccaaggtcat**-3′ |
| Reverse Transcription Adapter-oligoA | 5′-[5PHOS]-ggcttcttcttgctcttaggtagtaggttc-3′ | |

a) Specific RTA sequence is labeled in cursive

b) The 3'ends of the positive and negative RNA strands of each rotavirus genome segments sequence are labeled in bold.
